# Evolution of Virulence in Emerging Epidemics: From Theory to Experimental Evolution and Back

**DOI:** 10.1101/2024.03.13.584824

**Authors:** Wakinyan Benhamou, François Blanquart, Marc Choisy, Thomas W. Berngruber, Rémi Choquet, Sylvain Gandon

**Author notes:** These authors contributed equally to this work (Wakinyan Benhamou, François Blanquart). These authors also contributed equally to this work (Rémi Choquet, Sylvain Gandon).

## Abstract

The experimental validation of theoretical predictions is a crucial step in demonstrating the predictive power of a model. While quantitative validations are common in infectious diseases epidemiology, experimental microbiology primarily focuses on the evaluation of a qualitative match between model predictions and experiments. In this study, we develop a method to deepen the quantitative validation process with a polymorphic viral population. We analyse the data from an experiment carried out to monitor the evolution of the temperate bacteriophage λ spreading in continuous cultures of *Escherichia coli*. This experimental work confirmed the influence of the epidemiological dynamics on the evolution of transmission and virulence of the virus. A variant with larger propensity to lyse bacterial cells was favoured in emerging epidemics (when the density of susceptible cells was large), but counter-selected when most cells were infected. Although this approach qualitatively validated an important theoretical prediction, no attempt was made to fit the model to the data nor to further develop the model to improve the goodness of fit. Here, we show how theoretical analysis – including calculations of the selection gradients – and model fitting can be used to estimate key parameters of the phage life cycle and yield new insights on the evolutionary epidemiology of the phage λ. First, we show that modelling explicitly the infected bacterial cells which will eventually be lysed improves the fit of the transient dynamics of the model to the data. Second, we carry out a theoretical analysis that yields useful approximations that capture at the onset and at the end of an epidemic the effects of epidemiological dynamics on selection and differentiation across distinct life stages of the virus. Finally, we estimate key phenotypic traits characterizing the two strains of the virus used in our experiment such as the rates of prophage reactivation or the probabilities of lysogenization. This study illustrates the synergy between experimental, theoretical and statistical approaches; and especially how interpreting the temporal variation in the selection gradient and the differentiation across distinct life stages of a novel variant is a powerful tool to elucidate the evolutionary epidemiology of emerging infectious diseases.

## 1 Introduction

Evolutionary epidemiology theory predicts that the evolution of pathogen transmission is driven by the availability of susceptible hosts. At the onset of an epidemic, when the density of susceptible hosts is high, more transmission is favoured by natural selection. When a positive covariance exists between transmission and virulence (Alizon et al., 2009; Alizon & Michalakis, 2015; Anderson & May, 1982), this selection for higher transmission can indirectly select for higher virulence (Bull, 1994; Day, 2002; Day & Proulx, 2004; Frank, 1996; Gandon & Day, 2007; Lenski & May, 1994). Yet, an experimental validation of this prediction was needed to demonstrate the relevance of these predictions on the evolution of pathogens in emerging epidemics.

This prediction was put to the test in a previous study using experimental evolution of the temperate bacteriophage (or phage) λ (Berngruber et al., 2013). Phages are viruses that infect bacteria and phage λ is the archetypal temperate phage, which can switch between a lytic and a lysogenic life style. Upon infection, the virus may commit to the lytic pathway by hijacking the host’s replication machinery to produce new virions (viral particles) and eventually release them in the environment after the lysis of the host cell. Alternatively, the virus may commit to the lysogenic pathway by integrating its genome into the bacterial chromosome where it will lie in a dormant state as a prophage and be hereditarily transmitted to daughter cells at the pace of lysogen divisions. The prophage may also regain virulence by excising itself from the genome, switching to a lytic cycle (reactivation, also called induction) and thus shifting from vertical to horizontal transmission (Echols, 1972; Gandon, 2016; Lwoff, 1953; Ptashne, 1992). The evolution experiment designed in Berngruber et al., 2013 monitored the competition between two strains of phage λ with distinct life-history strategies in continuous cultures of *E. coli*. The first strain is the wildtype, which is known to have a relatively large lysogenization rate and low reactivation rate. The second strain is the λcI857 variant, which carries a point mutation in the transcriptional repressor protein cI (St-Pierre & Endy, 2008; Sussman & Jacob, 1962); the cI mutant is known to be more virulent and transmitted mostly horizontally through lytic cycles. Berngruber et al., 2013 developed a mathematical model tailored to the life cycle of phage λ. Numerical simulations of this model using parameter values from previous experimental studies led to three theoretical predictions: (i) the virulent strain outcompetes the wildtype when susceptible hosts are abundant, but the direction of selection is reversed as soon as the epidemic reaches high prevalence, (ii) the lower the initial prevalence, the higher the increase in virulence during the epidemic and (iii) the virulent strain is always more frequent among viral particles than among prophages. Tracking both the epidemiology (prevalence) and the evolution of the virus (frequency of the virulent strain among viral particles and among infected bacteria), all three predictions were confirmed experimentally (Berngruber et al., 2013). Yet, the data were only used as a qualitative validation of the theory and no attempt was made to explore the quantitative match between the predicted and the observed dynamics of the virus.

In the present work, we show how the quantitative analysis of the experimental results from Berngruber et al., 2013 improves our understanding of the evolutionary epidemiology of phage λ. First, we modified the structure of the epidemiological model to better capture the transient evolutionary dynamics of the virus among infected bacteria. Second, we carry out a theoretical analysis of this model to provide useful approximations to predict the evolutionary dynamics of the virus at different stages of the epidemic. In particular, we compute the selection gradients and the differentiation across distinct life stages of the virus. Finally, we develop a statistical inference approach to obtain quantitative estimates of the parameters of the model and, especially, the life-history traits of the different strains of phage λ.

## 2 Materials and methods

Following Berngruber et al., 2013, we first model the competition between the wildtype strain – hereafter denoted by the subscript *w* – and the mutant strain λcI857 – hereafter denoted by the subscript *m* – of phage λ in a chemostat with a well-mixed continuous culture of its bacterial host *E. coli* (**Fig. 1**). We summarize the notations in **Table 1**. We then recall the experimental data generated in the original study and describe how we generate simulated data to validate our ability to estimate model parameters. In the last section, we detail the statistical inference approach we developed.

**Table 1:**
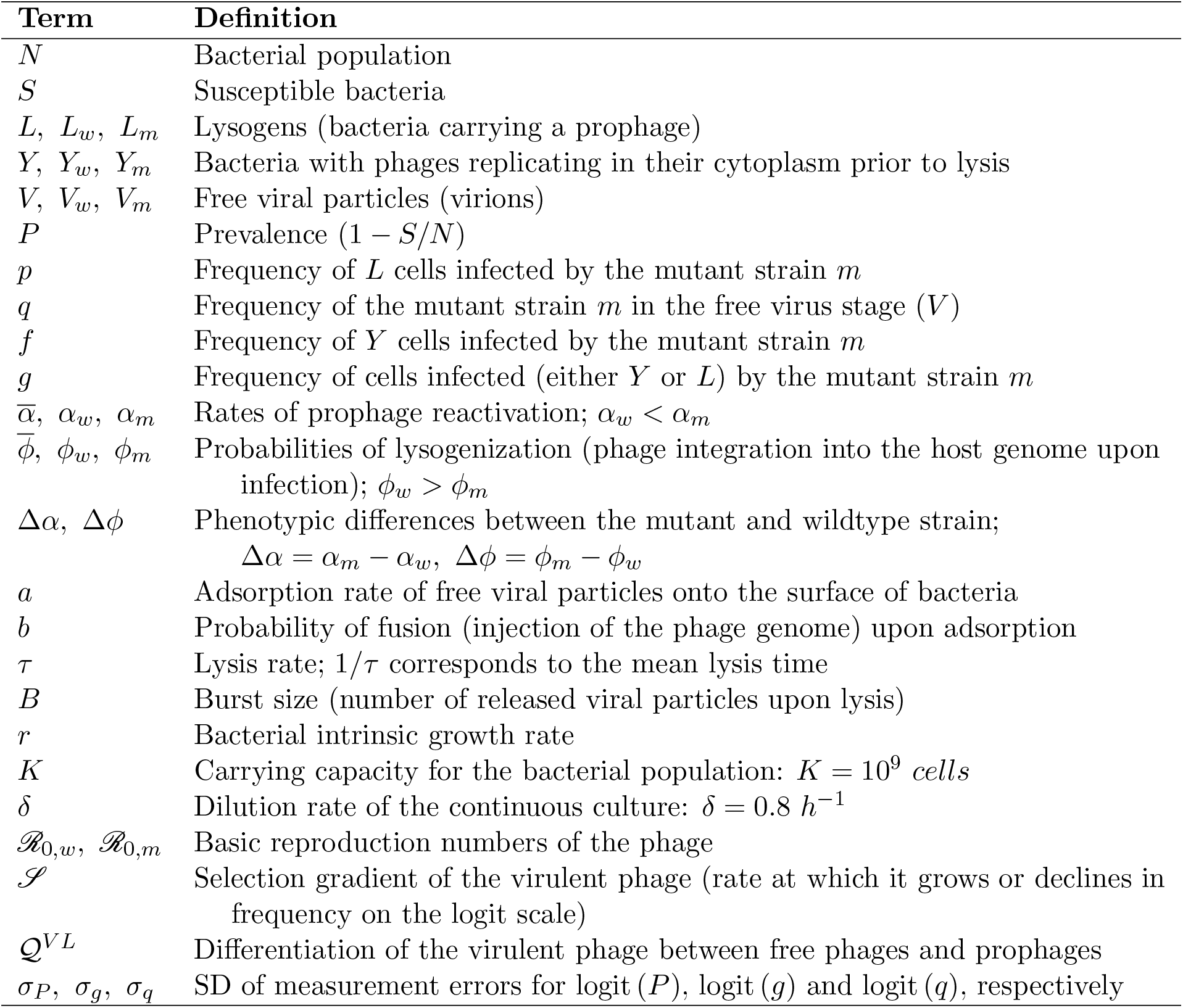
Notations. The subscripts *w* and *m* refer to the wildtype strain and the mutant (or virulent) strain λcI857 of phage λ, respectively. Overlines refer to mean values of life-history traits across all genotypes. SD stands for ‘standard deviation’.

| Term | Definition |
| --- | --- |
| $N$ | Bacterial population |
| $S$ | Susceptible bacteria |
| $L, L_w, L_m$ | Lysogens (bacteria carrying a prophage) |
| $Y, Y_w, Y_m$ | Bacteria with phages replicating in their cytoplasm prior to lysis |
| $V, V_w, V_m$ | Free viral particles (virions) |
| $P$ | Prevalence ($1 - S/N$) |
| $p$ | Frequency of $L$ cells infected by the mutant strain $m$ |
| $q$ | Frequency of the mutant strain $m$ in the free virus stage ($V$) |
| $f$ | Frequency of $Y$ cells infected by the mutant strain $m$ |
| $g$ | Frequency of cells infected (either $Y$ or $L$) by the mutant strain $m$ |
| $\bar{\alpha}, \alpha_w, \alpha_m$ | Rates of prophage reactivation; $\alpha_w < \alpha_m$ |
| $\bar{\phi}, \phi_w, \phi_m$ | Probabilities of lysogenization (phage integration into the host genome upon infection); $\phi_w > \phi_m$ |
| $\Delta\alpha, \Delta\phi$ | Phenotypic differences between the mutant and wildtype strain;<br>$\Delta\alpha = \alpha_m - \alpha_w, \Delta\phi = \phi_m - \phi_w$ |
| $a$ | Adsorption rate of free viral particles onto the surface of bacteria |
| $b$ | Probability of fusion (injection of the phage genome) upon adsorption |
| $\tau$ | Lysis rate; $1/\tau$ corresponds to the mean lysis time |
| $B$ | Burst size (number of released viral particles upon lysis) |
| $r$ | Bacterial intrinsic growth rate |
| $K$ | Carrying capacity for the bacterial population: $K = 10^9$ cells |
| $\delta$ | Dilution rate of the continuous culture: $\delta = 0.8 \text{ h}^{-1}$ |
| $\mathcal{R}_{0,w}, \mathcal{R}_{0,m}$ | Basic reproduction numbers of the phage |
| $\mathcal{S}$ | Selection gradient of the virulent phage (rate at which it grows or declines in frequency on the logit scale) |
| $\mathcal{Q}^{VL}$ | Differentiation of the virulent phage between free phages and prophages |
| $\sigma_P, \sigma_g, \sigma_q$ | SD of measurement errors for logit ($P$), logit ($g$) and logit ($q$), respectively |

**Figure 1:**
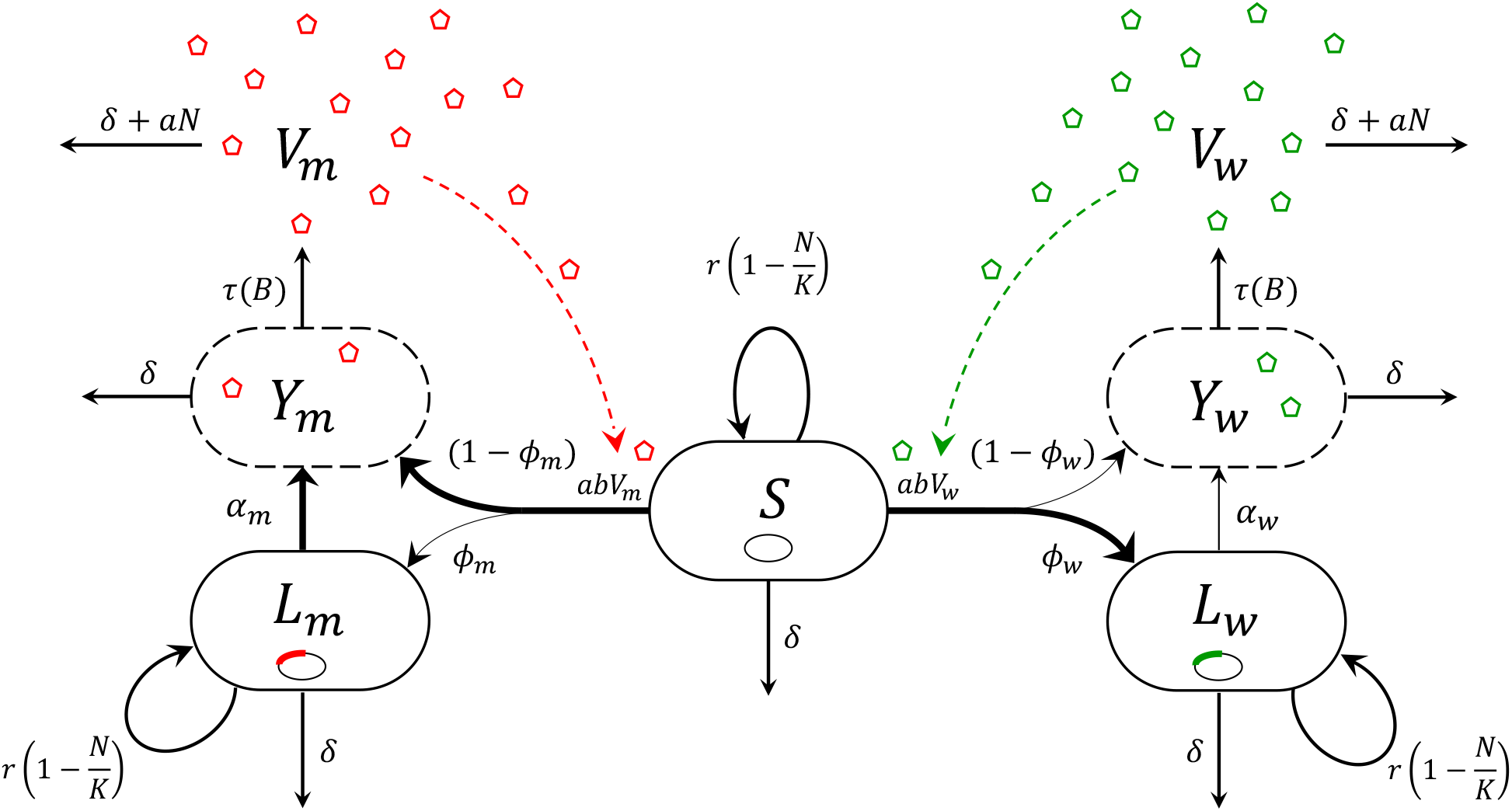
Flow chart of the phage-bacteria system. The subscripts *w* and *m* refer to the wildtype (in green) and mutant (or virulent, in red) strain of phage λ, respectively. In the bacterial population (*E. coli*) of size *N*, bacteria are either susceptible (*S*), lysogenic (*L*) – i.e., carrying a prophage in their chromosome (small ellipses) – or carrying phages (small pentagons) replicating in their cytoplasm prior to lysis (*Y*); *V* is the free virus stage (culture medium). Width of the arrows between *S, L* and *Y* reflects the relative rates of different events: the wildtype strain is transmitted mostly vertically (high probability of lysogenization and low rate of prophage reactivation) while the virulent strain is transmitted mostly horizontally (low probability of lysogenization and high rate of prophage reactivation). Dashed arrows symbolize the role of the free viral particles in the force of infection (epidemiological feedback). See notations in **Table 1**.

### 2.1 A model coupling epidemiology and evolution of phage virulence

#### 2.1.1 Epidemiology

Bacteria are either susceptible (*S*) or infected with phage λ (we do not consider resistant bacteria); infected bacteria may be lysogenic (*L*), following phage integration (lysogenization), or carrying phages replicating in their cytoplasm prior to lysis (*Y*), following either lytic infection or prophage reactivation. Unlike the original model in Berngruber et al., 2013, adding an extra stage *Y* allows us to take the lysis time into account; we show below that this new model improves the goodness of fit to the data (see **Results §3.2.2**). The free virus stage (*V*) corresponds to the free viral particles (virions) that are in the culture medium – “free” meaning “extracellular” here. Throughout, for each state variable (aka compartment), for instance *S*, we denote by *S*(*t*) its density at the current time *t* and 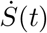 its derivative with respect to time. Hence, *L*(*t*) = *L*_*w*_(*t*) + *L*_*m*_(*t*) is the total density at time *t* of *L* cells, *Y* (*t*) = *Y*_*w*_(*t*) + *Y*_*m*_(*t*), of *Y* cells, *V* (*t*) = *V*_*w*_(*t*) + *V*_*m*_(*t*), of free viral particles, and *N* (*t*) = *S*(*t*) + *L*(*t*) + *Y* (*t*), of bacteria.

Susceptible and lysogenic bacteria grow at a *per capita* logistic rate *r*(1 − *N* (*t*)*/K*), where *r* is the intrinsic growth rate and *K* the carrying capacity. We assume, as in Berngruber et al., 2013, that the prophage does not affect the intrinsic growth rate of its host and that vertical transmission is perfect. Bacteria and virions are removed from the chemostat at a dilution rate *δ*. Free viral particles adsorb onto bacterial cells at a rate *a*; adsorption is non-reversible, that is, the fate of adsorbed viruses is only to infect or die. Infection of a susceptible host also requires the injection of the phage’s genetic material into the bacterial cytoplasm (fusion, with probability *b*). The force of infection – i.e., the *per capita* infection rate – is therefore given by *abV* (*t*). We assume that superinfection (including co-infection with both strains) does not occur. In particular, prophage establishment of phage λ is known to provide cellular immunity, or superinfection inhibition (Berngruber et al., 2010; Gandon, 2016; Lwoff, 1953; Ptashne, 1992). Upon infection, the wildtype and virulent strains of phage λ may either be integrated as prophages into the host genome (lysogenic cycle) with probabilities *ϕ*_*w*_ and *ϕ*_*m*_, respectively, such that Δ*ϕ* = *ϕ*_*m*_ −*ϕ*_*w*_ < 0, or start the biosynthesis and assembly of viral copies (lytic cycle) with complementary probabilities. Once integrated, reactivations of the wildtype and virulent prophages occur at a rate *α*_*w*_ and *α*_*m*_, respectively, such that Δ*α* = *α*_*m*_ − *α*_*w*_ > 0. Following either lytic infections or prophage reactivations, host cells are lysed at a rate *τ* and eventually release *B* viral particles upon lysis (burst size). We assume that parameters *a, b, τ* and *B* are the same for the wildtype and the mutant which yields the following system of ordinary differential equations (ODEs):

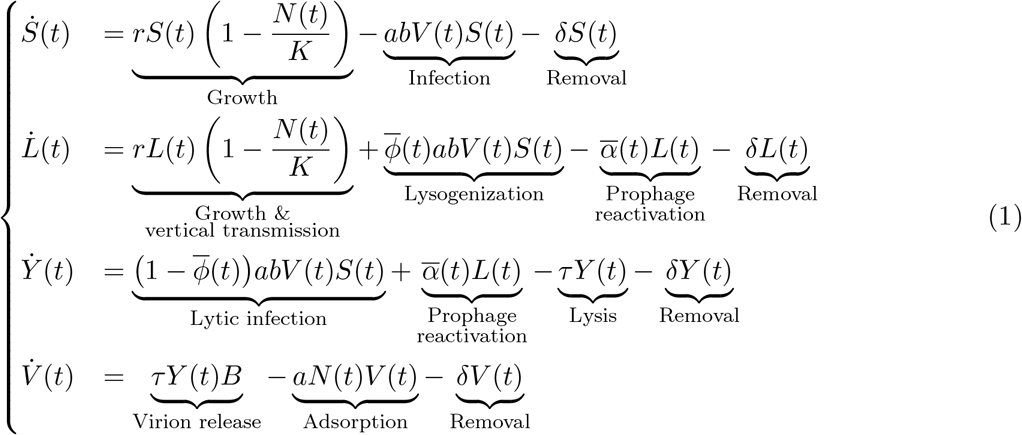

where 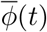 is the mean probability of lysogenization upon infection and 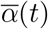 the mean rate of reactivation among prophages:

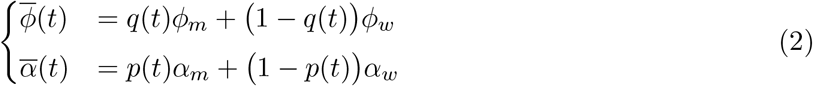

with *q*(*t*) = *V*_*m*_(*t*)*/V* (*t*) and *p*(*t*) = *L*_*m*_(*t*)*/L*(*t*), the frequencies of the virulent strain at time *t* among compartments *V* and *L*, respectively.

#### 2.1.2 Evolution

Along with these epidemiological dynamics, we also track the evolutionary dynamics of phage λ. We recall that *p*(*t*) and *q*(*t*) refer to frequencies of the virulent strain in compartment *L* and *V*, respectively. In addition, we also denote *f* (*t*) = *Y*_*m*_(*t*)*/Y* (*t*), the frequency of the virulent strain at time *t* among *Y* cells, and *g*(*t*), the frequency of the virulent strain in infected cells (either *Y* or *L*) such that:

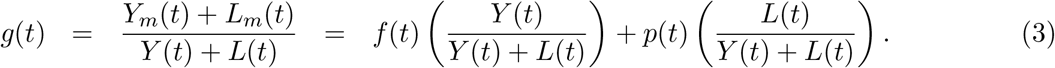

Using (1) and (2), one may calculate the ODE for each of these frequencies (see **SI Appendix §S1.2**). More conveniently, we will then focus on logit-frequencies instead, that is, the log odds ln (frequency of the mutant / frequency of the wildtype). Taken together, the equations of the temporal dynamics of the frequencies and the model (1) yield the coupled evolutionary-epidemiological dynamics of this phage-bacteria system. The analysis of this model can provide key insight on the evolutionary forces acting on the virus; it may also provide useful approximations for the change in mutant frequency at different stages of the epidemic.

### 2.2 Time series datasets

#### 2.2.1 Experimental data

We use experimental time series obtained in the first evolution experiment of Berngruber et al., 2013. Briefly, this experiment started with eight independent chemostats (5 mL chamber volume, maintained at 35°C) of well-mixed bacterial cultures of *E. coli* MG1655 (RecA+) at carrying capacity. Bacteria were initially infected by both the wildtype and virulent strain λcI857 of phage λ (prophage stage with initial ratio 1:1). Two treatments were considered using 4 chemostats each: (i) an epidemic treatment – low initial prevalence, around 1% – and (ii) an endemic treatment – high initial prevalence, around 99%. Throughout the course of the experiment, several quantities were monitored: the prevalence, the frequency infected by each strain and the strain frequency in the culture medium (free virus stage). Samplings in each chemostat were performed hourly, from 1 to 60 h maximum. The prevalence and the frequency of infected hosts were tracked using flow cytometry (FACS) with fluorescent protein marker colors (CFP and YFP) while the strain frequency in the free virus stage was tracked by qPCR.

#### 2.2.2 Simulated data

Alongside experimental data, we also carry out an analysis based on simulated data in order to validate our ability to infer parameters from experimental data. For this purpose, we take: *α*_*w*_ = 7 × 10^−3^, *α*_*m*_ = 2 × 10^−2^, *ϕ*_*w*_ = 0.2, *ϕ*_*m*_ = 2 × 10^−2^, *a* = 3 × 10^−9^, *b* = 0.1, *B* = 80, *r* = 1.4, *τ* = 1.5, *K* = 10^9^ and *δ* = 0.8. At *t* = 0, bacteria are at carrying capacity *K* with initial prevalence 1% (epidemic treatment) or 99% (endemic treatment) and the initial prophage ratio for the two strains is 1:1. We simulate the deterministic model (1) for the epidemic and endemic treatment (**Fig. S1**). We then add i.i.d. Gaussian noise at each time point to mimic measurement errors on the logit-prevalence logit (*P*(*t*)), the logit-frequency of hosts infected by the virulent phage logit(*g*(*t*)) and the logit-frequency of the virulent phage in the free virus stage logit (*q*(*t*)) – this is independently repeated four times for each simulation to obtain four replicates (chemostats) per treatment. We modulate data quantity through two sampling frequencies: 0.1 *h*^−1^ vs. 1 *h*^−1^, and we modulate data quality through two standard deviations (SD) of measurement errors: 0.01 vs. 0.5. We thus end up with four combinations of data quantity and quality (see example in **Fig. S2**) from which we then try to recover parameter values.

### 2.3 Maximum likelihood estimation

For the estimation process, we used a two-step approach. Using theoretical results, we first compute point estimates of the rates of prophage reactivation *α*_*w*_ and *α*_*m*_. Then, we fix the latter to estimate the remaining parameters of the model using non-linear optimizations. Note that it is possible to run non-linear optimizations to estimate all parameters but this two-step approach makes the optimization easier by reducing the dimensionality of the problem. The Sieve bootstrap method (Bühlmann, 1997; Ulloa et al., 2013) is used to compute the joint distributions of all these estimated parameters.

#### 2.3.1 Estimation of the rates of prophage reactivation

From the analysis of the model (see **Results §3.1**), we show that when the system reaches high prevalence the selection gradient ℐ of the mutant – i.e., the rate at which it grows or declines in frequency on the logit scale – is simply given by: ℐ = *α*_*w*_ − *α*_*m*_ = −Δ*α* (**Results §3.1.2**). Furthermore, the differentiation 𝒬^*V L*^ of the virulent strain between free phages and prophages converges towards approximately: 𝒬^*V L*^ = *α*_*m*_*/α*_*w*_ = 1 + Δ*α/α*_*w*_ (**Results §3.1.3**). Combining these two expressions enables us to estimate separately both reactivation rates:

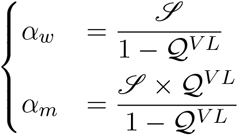

For each chemostat, we therefore only keep the data from the time point the prevalence has reached 95%. We fit a linear model on the logit-frequency infected by the virulent phage logit (*g*(*t*)) to estimate the slope ℐ, and we estimate 𝒬^*V L*^ by calculating the geometric mean of 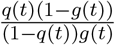 (see details in **SI Appendix §S3.1**). Note that we substitute *p*(*t*) by *g*(*t*) for the calculation of the differentiation because we only have access to *g*(*t*) in the experiment, and *g*(*t*) is almost identical to *p*(*t*) towards the end of the epidemic (see **Results §3.1.2** and **Fig. S1**).

#### 2.3.2 Estimation of the remaining parameters

We have three response variables: (i) logit (*P* (*t*)), the logit-prevalence, (ii) logit (*g*(*t*)), the logit-frequency of hosts infected by the virulent phage and (iii) logit(*q*(*t*)), the logit-frequency of the virulent phage in the free virus stage. Let 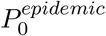 and 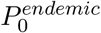 be the initial conditions (at *t* = 0) of the prevalence in the epidemic and endemic treatments, respectively, and *p*_0_, the initial condition of the frequency infected by the virulent phage (identical for both treatments). Let *θ* = (*ϕ*_*w*_, *ϕ*_*m*_, *a, b, r, τ*, 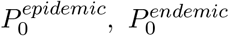, *p*_0_) be the vector of model parameters to estimate. Several parameter values are fixed: *K* = 10^9^ *cells* and *δ* = 0.8 *h*^−1^ (Berngruber et al., 2013); *α*_*w*_ and *α*_*m*_ are fixed to their previous point estimates (cf. **Methods §2.3.1**); and *B* = 80 *virus*. *cell*^−1^ (Wang, 2006) as we show in **Results §3.2.1** that the burst size *B* is not separately identifiable from parameter *b*. We estimate as well *σ*_*P*_, *σ*_*g*_ and *σ*_*q*_, the SD of the measurement errors for each of the three response variables, respectively. We assume that variation around the deterministic dynamics stems solely from measurement errors and we thereby neglect any additional process stochasticity. This is justified in particular by the very controlled conditions of the experiment and by the large population sizes of both the bacteria and the phage in the chemostats. For each response variable, measurement errors are assumed to be normally distributed and i.i.d. across all treatments, replicates and time points. The corresponding likelihoods (ℒ) are thence respectively given by:

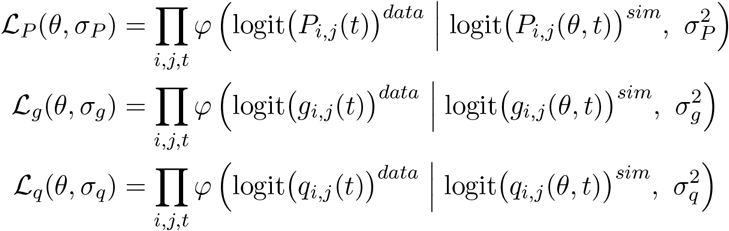

where *i* refers to treatments (epidemic vs. endemic), *j* to the *j*th replicate (chemostat), *t* to time points and *φ* (. | *µ, σ*^2^) to the probability density function of the Normal distribution with mean *µ* and variance *σ*^2^; *sim* indicates model outputs. We then denote 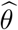, the MLE estimator of *θ*, such that:

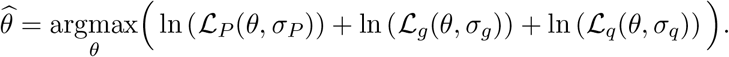

In practice, we minimize the negative overall log-likelihood using the Nelder-Mead (aka downhill simplex) algorithm (Nelder & Mead, 1965). Parameter bounds (reported in **Table S2**) are imposed through parameter transformations. Due to the presence of local minima, optimizations are repeated for 2,000 sets of uniformly drawn starting points to ensure convergence to a global minimum. The best MLE set of estimates 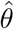 corresponds to the fit associated with the lowest negative overall log-likelihood with successful completion.

#### 2.3.3 Confidence intervals

We generate bootstrapped data to compute 95% CIs of our parameters using Sieve bootstrap (Bühlmann, 1997; Ulloa et al., 2013) on the residuals between experimental data and the best fit of our model. For this purpose, autoregressive moving-average (ARMA) models are fitted to the time series of centred residuals of each chemostat independently. We then use ARMA models to simulate new residuals from which we reconstruct new datasets. We eventually reiterate the above estimation procedure (cf. **§2.3.1-2.3.2**), but starting non-linear optimizations only from the best MLE estimates we obtained with the original data. By repeating this for 10,000 bootstrapped datasets, we compute the joint distributions of estimated parameters.

### 2.4 Details of the implementation

Numerical simulations and data analyses were carried out using R (R Core Team, 2022) version 4.2.0 (2022-04-22). ODEs were solved numerically by the function `ode` – with method `lsoda` – from the package `deSolve` (Soetaert et al., 2010). Non-linear optimizations for MLE were tackled with the function `nmk` (Kelley, 1999), from the package `dfoptim`, which gave here more stable results than `optim` from base R. Fit and selection of ARMA models for Sieve bootstrap were carried out using the function `auto.arima`, from the package ‘*forecast* ‘ (Hyndman & Khandakar, 2008).

## 3 Results

### 3.1 Theoretical analysis

When *r* > *δ*, the virus-free system converges to an equilibrium where *S*(∞) = *K*(1 − *δ/r*). When a single strain *k* ∈ {*w, m*} of the virus (with phenotypes *α*_*k*_ and *ϕ*_*k*_) is introduced in the bacterial population (fully susceptible, with density *S*_0_), the fate of the phage-bacteria system depends on the basic reproduction number of the pathogenℛ_0,*k*_ which is given by:

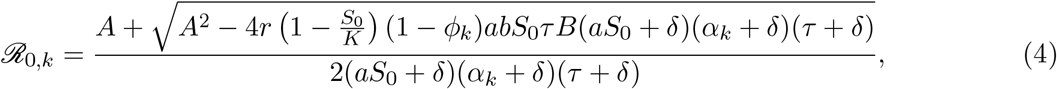

With 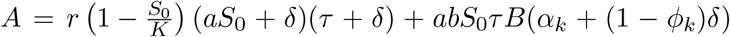 (see **SI Appendix §S2.1** for the construction of the next-generation matrix, following (Diekmann et al., 2010)). When ℛ_0,*k*_ < 1, the virus goes extinct and the bacterial population converges to the previous virus-free equilibrium. Alternatively, when ℛ_0,*k*_ > 1, an epidemic breaks out and eventually stabilises to an endemic equilibrium where all the cells are infected by the virus (**SI Appendix §S2.1**).

In the following, we analyse the evolutionary dynamics (i) at the beginning of an epidemic where *S*(*t*) = *S*_0_ and (ii) when the system stabilizes towards the endemic equilibrium where *S*(*t*) = 0.

#### 3.1.1 Evolution in an emerging epidemic

At the onset of the epidemic, susceptible cells are highly abundant. For the sake of simplicity, we analyse the dynamics of the epidemic when the host density is assumed to be constant over time (*S*(*t*) = *S*_0_). As in the experimental design, we start with a bacterial population at carrying capacity (*S*_0_ ≈ *K*). Density dependence reduces cell reproduction and vertical transmission of the virus. Consequently, the epidemic is mainly driven by the lytic pathway. At *t* = 0 though, only lysogens are introduced at very low density. After a very short time, the phage-bacteria system reaches its new dynamical regime, following prophage reactivations, lyses and releases of virions. From there, we assume that *L*(*t*)*/Y* (*t*) ≈ 0 and we focus on the dynamics of the mutant in the compartment *Y*. We derive an approximation of the selection gradient ℐ of the virulent phage, which corresponds to the rate at which it increases in frequency on the logit scale, and we show in **SI Appendix §2.2** that:

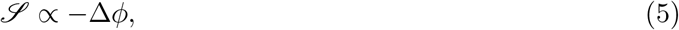

meaning that it is selected for (Δ*ϕ <* 0, cf. **Fig. S3-A**). This is consistent with the experimental data where the virulent phage transiently outcompetes the wildtype at the early stage of the epidemic treatment (Berngruber et al., 2013). Our prediction is accurate when the density of susceptible hosts remains effectively constant over time; otherwise, however, our prediction deviates from the simulation whose rate slows down because the density of susceptible cells is rapidly decreased by the spread of the epidemic (**Fig. 2-A** and **Fig. S3-C**).

**Figure 2:**
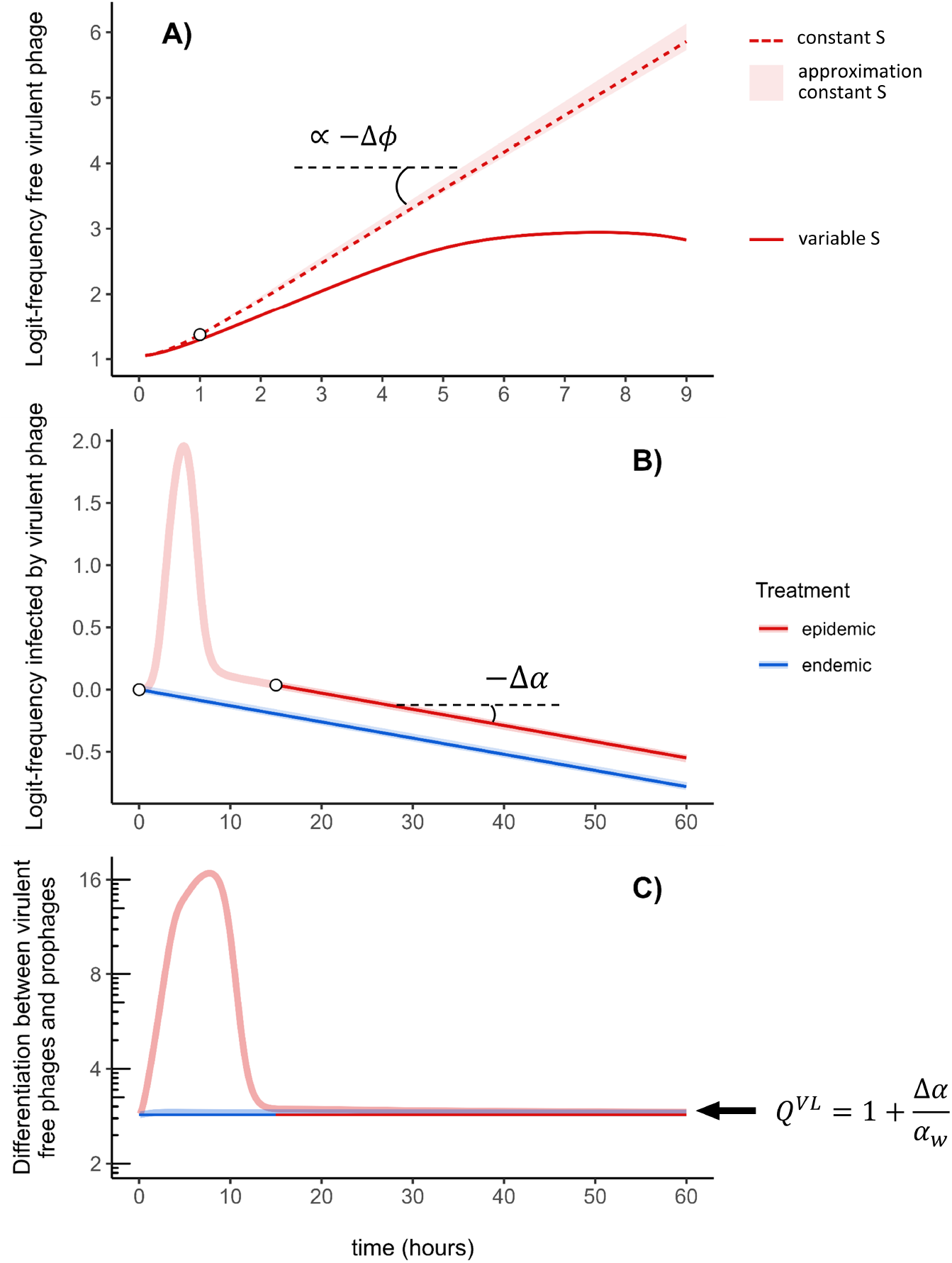
Theoretical predictions. At *t* = 0, bacteria are at carrying capacity *K* with initial prevalence 1% (epidemic treatment) or 99% (endemic treatment). The initial prophage ratio for the two strains is 1:1. See **Table S1** for parameter values. (A) The virulent phage is selected for at the early stage of the epidemic. Our approximation for the trajectory of the logit-frequency of the virulent phage (light red shading, see **SI Appendix §S2.2**) predicts the dynamics of the virulent phage when the density of susceptible cells is kept constant (dashed red line). The slope of the upper bound of our approximation is proportional to −Δ*ϕ* = *ϕ*_*w*_ − *ϕ*_*m*_ (equation (5)). As only lysogens are introduced in the susceptible bacterial population at *t* = 0, we let the phage-bacteria system reach its new dynamical regime and only start our approximation at *t* = 1 (white dot). However, the rapid drop in the availability of susceptible hosts due to the viral epidemic reduces the increase of the virulent phage relative to our approximation (compare the full red line and the dashed red line). (B) The virulent phage is counter-selected when the epidemic reaches high prevalence (indicated with the white dots in the epidemic and endemic treatments). At the end of the epidemic, the logit-frequency of the virulent phages decreases linearly with negative slope −Δ*α* = *α*_*w*_ − *α*_*m*_ (equation (6)). (C) The virulent strain is more frequent among free viruses (*V*) than among prophages (*L*). When the system reaches high prevalence, we predict the differentiation between these two compartments converges towards an equilibrium that is approximately equal to 1 + Δ*α/α*_*w*_ = *α*_*m*_*/α*_*w*_ (equation (8)).

#### 3.1.2 Evolution at the end of the epidemic

At the end of the epidemic, we expect that all the cells will be infected by a prophage and there will no longer be any susceptible cells. Consequently, no horizontal transmission takes place and, in contrast with the previous scenario, we can neglect the density of *Y* cells relative to the density of lysogens. Indeed, since 1*/α*_*w*_ ≫ 1*/τ* (time elapsed between phage integration and reactivation is much longer than lysis time), then *Y* (*t*)*/L*(*t*) ≈ 0 (see details in **SI Appendix §S2.3**). Note that this also means that the frequency infected by the virulent strain is mainly driven by the corresponding frequency in *L*, that is, *g*(*t*) ≈ *p*(*t*). We show in **SI Appendix §S2.3** that the selection gradient of the virulent phage is given by:

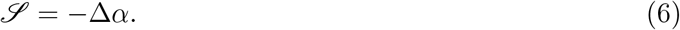

As a result, the virulent phage is counter-selected in the long-term (Δ*α* > 0) and, in each compartment, linearly declines in frequency at a rate ℐ (negative slope) on the logit scale (**Fig. 2-B** and **Fig. S1-B**).

#### 3.1.3 Differentiation across compartments

The two previous subsections deal with the dynamics of evolution over time; let us now look at the dynamics of selection between different compartments. We define here the differentiation 𝒬^*VL*^ between the free virus stage and lysogens as:

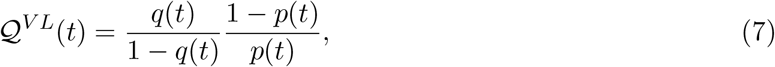

such that:

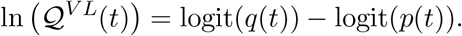

We show in **SI Appendix §S2.4** that, at the end of the epidemic, the differentiation 𝒬^*V L*^(*t*) converges approximately towards:

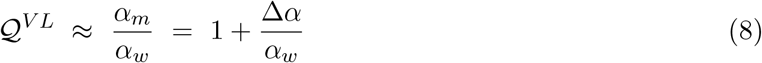

(**Fig. 2-C**), a value of 1 corresponding to no differentiation. We obtain the same expression for the convergence of the differentiation between *Y* and *L* cells and around 1 between the free virus stage *V* and *Y* cells (cf. **SI Appendix §2.4**). Interestingly, these results are also valid at *t* = 0^+^ (**Fig. S4**) as there are no free phage particles yet and thereby no horizontal transmission which, within a very short space of time, resembles the end of the epidemic. When the system stabilizes at the end of the epidemic, the virulent strain is therefore more frequent among free viruses and *Y* cells than among *L* cells (prophages) but we also expect almost no differentiation between free viruses and *Y* cells (**Fig. S4**).

### 3.2 Statistical inference

#### 3.2.1 Inference from simulated data

We first conduct a simulation study. We start by validating our estimation method of the rates of prophage reactivation *α*_*w*_ and *α*_*m*_. Combining equations (6) and (8), we compute point estimates of both parameters (cf. **Methods §2.3.1**) for a large number of simulated datasets. Overall, estimated values show a very good match with those used in the original simulations (**Fig. S5**), especially when the SD of measurement errors is small.

We then show that parameters *b* and *B* may not be separately identifiable using the simulated dataset closest to the original deterministic simulation (sampling effort = 10 *h*^−1^ and SD of measurement error = 0.01, see **Fig. S2-A**). We fix *K* and *δ* to their known values, as well as *α*_*w*_ and *α*_*m*_, as though the rates of prophage reactivation have been correctly estimated beforehand. Setting (*b, B*) to many combinations of values ((*b, B*) ∈ [0, 0.2] × [0, 100]) and maximizing over the remaining parameters, the resulting landscape of the overall log-likelihood suggests that only the product *b × B* is identifiable (**Fig. S6**). Such compensatory effect may be understood as the impossibility here of discriminating whether phage λ infect fewer bacteria (lower probability *b* of fusion) but release more viral copies upon lysis (larger burst size *B*) or vice-versa. This is the reason why we fix *B* in the following and only estimate *b*.

Fixing *B* to its true value used in the original simulations, point estimates of parameters are quite close to their true values (**Fig. S7**). However, some parameters – like *r, τ* or *b* – seem more difficult to estimate with a lower sampling effort.

#### 3.2.2 Inference from experimental data

The original model used in Berngruber et al., 2013 failed to properly capture the evolutionary dynamics among infected hosts at the early stage of the epidemic. Such discrepancy was an opportunity to go back from experimental data to theory and to challenge the structure of the model. We noticed that adding an extra stage *Y* in the infected compartment allowed us to better fit the experimental data than the original model (ΔAIC=-101). This significant improvement strongly supported the new version of the model developed in this study and for which we present the estimation results below.

First, we estimate: Δ*α* = 9.31 × 10^−3^ *h*^−1^ and 𝒬^*V L*^ = 4.60 (see **Fig. 3** for the latter). Point estimates of the rates of prophage reactivation (expressed in *h*^−1^) are thus: 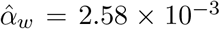 and 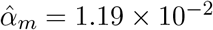

Second, we perform non-linear optimizations to estimate the remaining parameters. Overall, computed dynamics of the logit-prevalence and logit-frequencies of the virulent phage closely fit experimental data (**Fig. 4**). Best MLE estimates are listed in **Table 2**, along with their 95% bootstrap-based CIs (see **Fig. S8** for the density distributions of estimated parameters). We show pairwise correlation coefficients in **Fig. S9**. Some parameters are correlated positively, in particular: *α*_*w*_ with *α*_*m*_, *p*_0_ with both reactivation rates, *ϕ*_*m*_ with 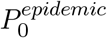 and *r* with *b*; or negatively, in particular: *τ* with *b* and *r*. In **Fig. S10**, we plot the 95% distributions of fitted values. To test the impact of the fixed value used for the burst size *B*, we study how perturbations of the original value (80 *virus*. *cell*^−1^) affect the estimation of the other parameter values (new point estimates from non-linear optimizations). In **Fig. S11**, we show that all parameters are extremely robust to these perturbations, except for parameter *b* as expected from **Fig. S6** (but the product *b × B* is always around 3.95). It is also worth noting that the basic reproduction number is not affected by the choice of the value of *B*, as it always appears as a product with *b* in (4). We estimate the basic reproduction number of the wildtype strain *ℛ*_0,*w*_ to be 1.48 (95% CI [1.04, 2.15]) and of the virulent strain *ℛ*_0,*m*_ to be 2.20 (95% CI [1.58, 3.05]). Interestingly, the basic reproduction number of the virulent mutant *ℛ*_0,*m*_ is higher than that of the wildtype *ℛ*_0,*w*_, but the virulent mutant always loses the competition in the long term. The basic reproduction number gives the expected number of secondary infections in an otherwise fully susceptible host population (Anderson & May, 1991). The generation interval distribution of the two strains being similar for the lytic cycle, the basic reproduction number provides a good predictor of the relative fitness of the two strains at the early stage of the epidemic (Park et al., 2019; Wallinga & Lipsitch, 2007): the mutant outcompetes the wildtype as *ℛ*_0,*m*_ > *ℛ*_0,*w*_. When the density of susceptible host cells drops, however, the basic reproduction number becomes a poor predictor of fitness. Eventually, only the strain that tolerates the worst environment in terms of resource density – in this case, the lowest density of susceptible host cells – survives (pessimization principle) (Diekmann, 2004). The wildtype replicates better at lower *S* cells densities than the virulent strain because it relies more on vertical transmission and less to transmission to new susceptible cells. *ℛ*_0_ is not maximized in the long-term because the “niche” of the virus is multidimensional, again because of the two modes of replication (Lion & Metz, 2018).

**Table 2:**
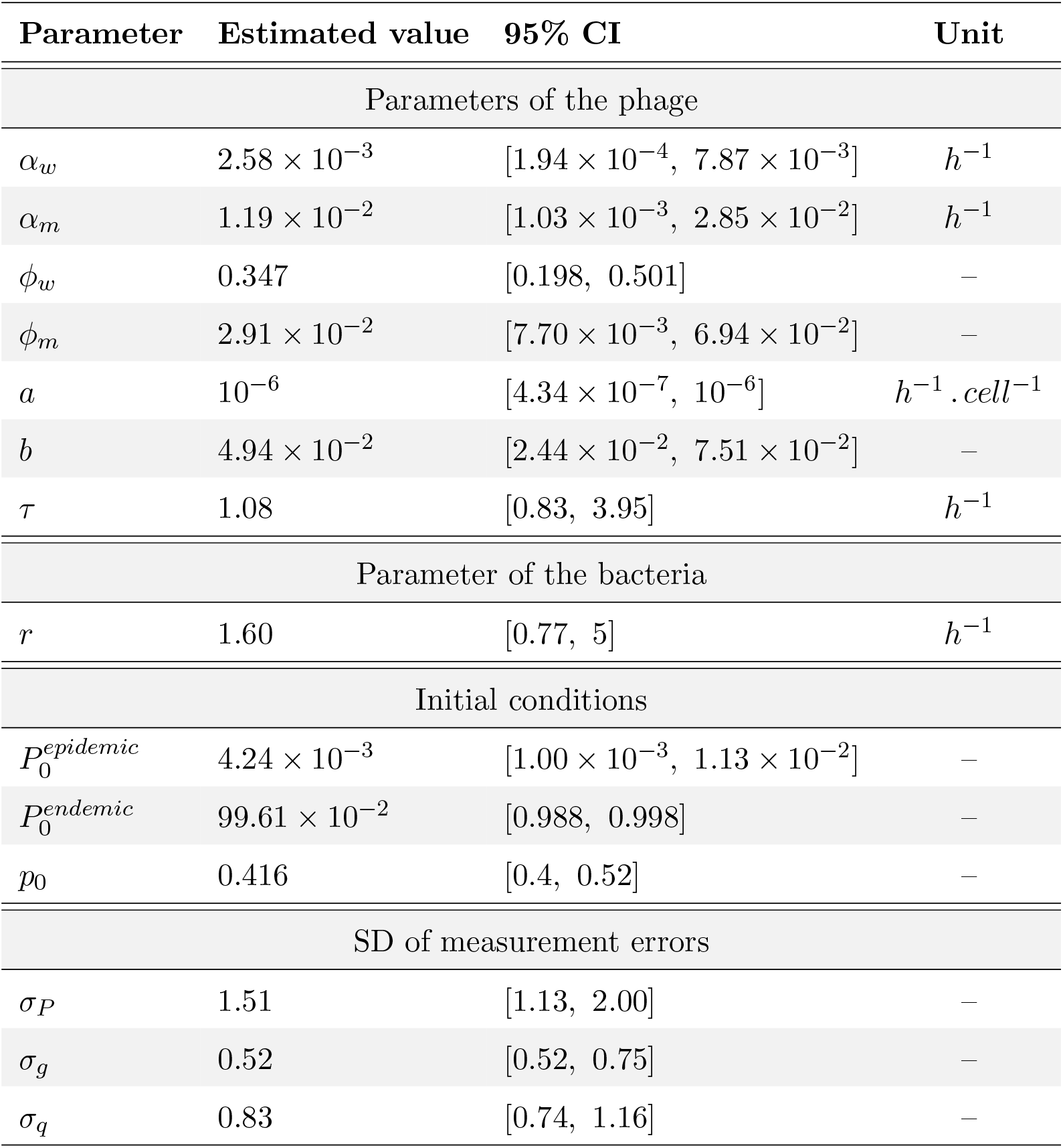
MLE parameter estimations. In a first step, parameters *α*_*w*_ and *α*_*m*_ are estimated directly using analytic approximations and linear models. In a second step, we fix *α*_*w*_ and *α*_*m*_ to their previous point estimates and, starting from 2,000 sets of initial values, non-linear optimizations maximizing the overall log-likelihood with the experimental data are run to estimate the remaining parameters. 95% CIs are based on Sieve bootstrap (6,686 sets of estimates). Throughout, fixed parameters are: *K* = 10^9^ *cells, δ* = 0.8 *h*^−1^ and *B* = 80 *virus*. *cell*^−1^.

| Parameter | Estimated value | 95% CI | Unit |
| --- | --- | --- | --- |
| Parameters of the phage |  |  |  |
| $\alpha_w$ | $2.58 \times 10^{-3}$ | $[1.94 \times 10^{-4}, 7.87 \times 10^{-3}]$ | $h^{-1}$ |
| $\alpha_m$ | $1.19 \times 10^{-2}$ | $[1.03 \times 10^{-3}, 2.85 \times 10^{-2}]$ | $h^{-1}$ |
| $\phi_w$ | 0.347 | [0.198, 0.501] | — |
| $\phi_m$ | $2.91 \times 10^{-2}$ | $[7.70 \times 10^{-3}, 6.94 \times 10^{-2}]$ | — |
| $a$ | $10^{-6}$ | $[4.34 \times 10^{-7}, 10^{-6}]$ | $h^{-1} \cdot cell^{-1}$ |
| $b$ | $4.94 \times 10^{-2}$ | $[2.44 \times 10^{-2}, 7.51 \times 10^{-2}]$ | — |
| $\tau$ | 1.08 | [0.83, 3.95] | $h^{-1}$ |
| Parameter of the bacteria |  |  |  |
| $r$ | 1.60 | [0.77, 5] | $h^{-1}$ |
| Initial conditions |  |  |  |
| $P_0^{epidemic}$ | $4.24 \times 10^{-3}$ | $[1.00 \times 10^{-3}, 1.13 \times 10^{-2}]$ | — |
| $P_0^{endemic}$ | $99.61 \times 10^{-2}$ | [0.988, 0.998] | — |
| $p_0$ | 0.416 | [0.4, 0.52] | — |
| SD of measurement errors |  |  |  |
| $\sigma_P$ | 1.51 | [1.13, 2.00] | — |
| $\sigma_g$ | 0.52 | [0.52, 0.75] | — |
| $\sigma_q$ | 0.83 | [0.74, 1.16] | — |

**Figure 3:**
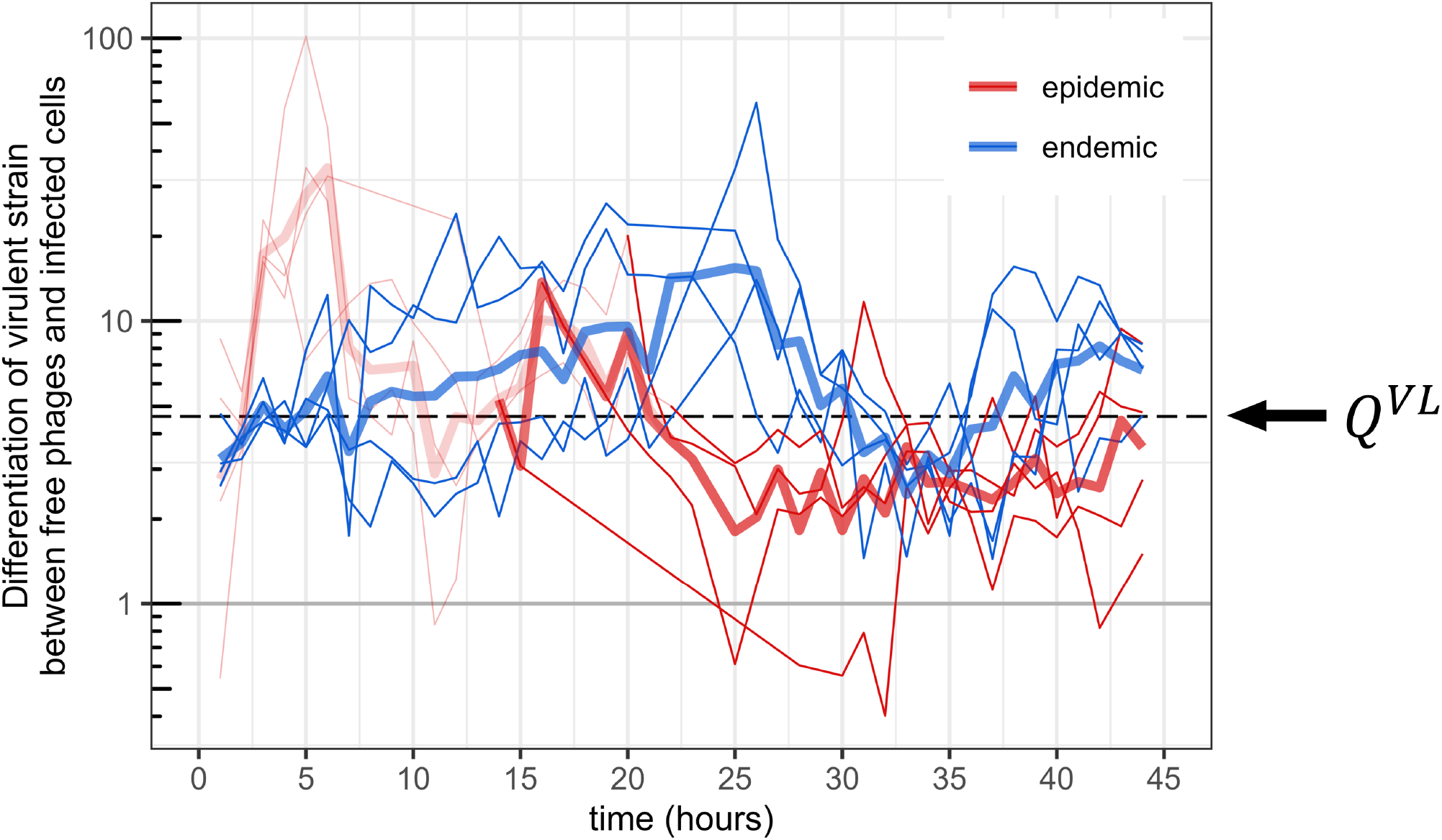
Differentiation of the virulent strain between free phages and infected cells. We compute *q*(*t*)(1 − *g*(*t*))*/* ((1 − *q*(*t*))*g*(*t*)) for each chemostat (thin lines) and the corresponding geometric mean for each treatment (thick lines). The value 1 (horizontal grey line) corresponds to no differentiation. For each chemostat, values before the time point the system has reached a prevalence of 95% are shown in transparent color. With high prevalence, we have *g*(*t*) ≈ *p*(*t*) and we thereby obtain the differentiation of the virulent strain between free phage particles (*V*) and prophages (*L*). In these conditions, such differentiation is expected to reach an equilibrium – the horizontal dashed line indicates the corresponding geometric mean (4.60) across all chemostats and time points – with theoretical value 1 + Δ*α/α*_*w*_ = *α*_*m*_*/α*_*w*_ (cf. **Fig. 2-C**).

**Figure 4:**
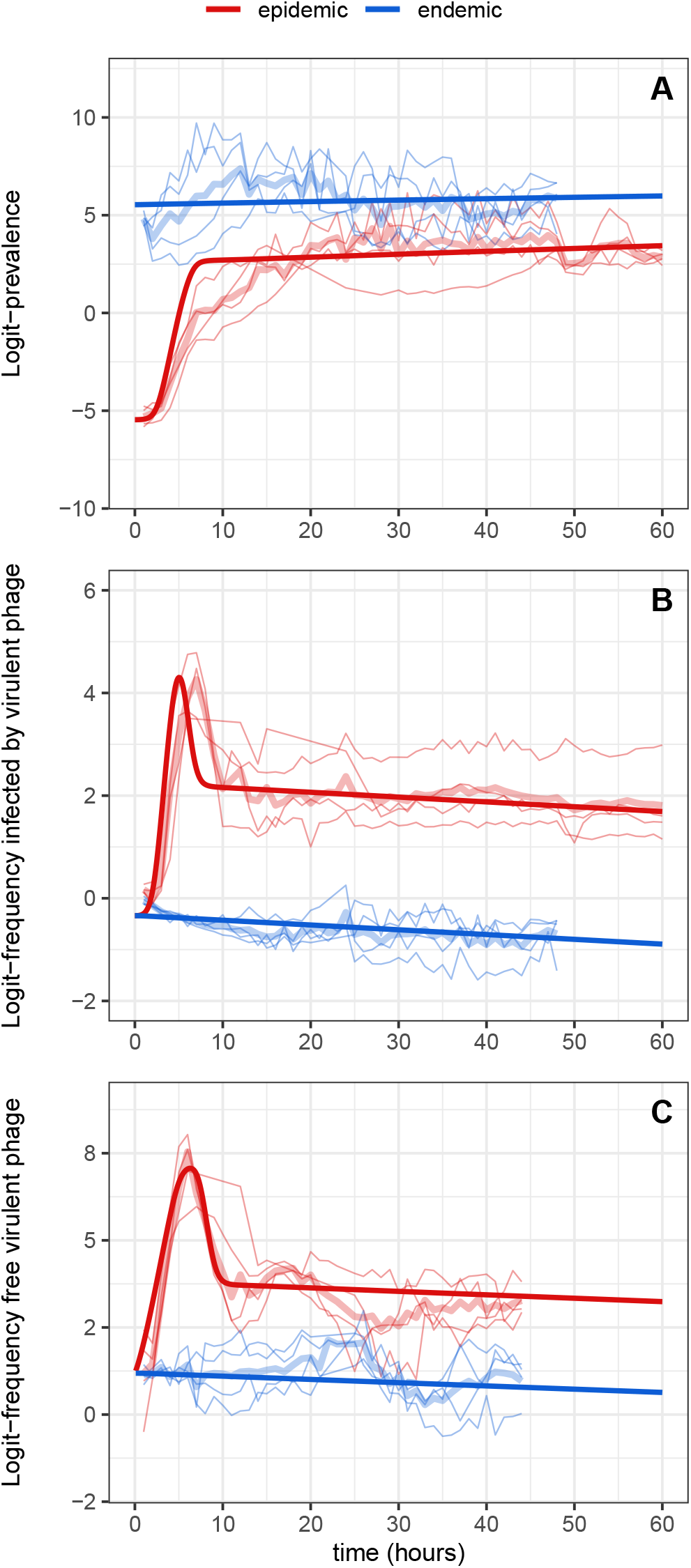
Fitted values on experimental data. Fitted values (thick dark lines) were obtained from the simulation based on the best MLE estimations (see **Table 2**). We estimate model parameters using experimental data (light-colored thin lines) from the evolution experiment designed in Berngruber et al., 2013; initial prevalence is either low (epidemic condition, in red) or high (endemic condition, in blue). Light-colored thick lines correspond to mean logit-values across all replicates per treatment. (A) Logit-prevalence of infection; (B) logit-frequency of cells infected (involved either in a lysogenic or a lytic cycle) by the the mutant (virulent) phage; (C) logit-frequency of the mutant (virulent) in the culture medium (free virus stage).

We compared our estimates of model parameters with previous studies (see reported values in **Table S3**), focusing in particular on the parameters of the phage. Note that previous estimations of these life-history traits were obtained from experimental assays using monomorphic phage cultures. Our estimate of the rate of prophage reactivation of the wildtype strain *α*_*w*_ falls in the range of previous *in vitro* estimates, which is very large (between 10^−7^ and 10^−2^ *h*^−1^ (De Paepe et al., 2016; Little et al., 1999; Zong et al., 2010)), probably due to the variability in experimental methods. For the cI variant, the orders of magnitude of *α*_*m*_ are more consistent (De Paepe et al., 2016; Zong et al., 2010). Other studies, using single-cell monitoring of infected cells to estimate the probability of lysogenization of the wildtype strain *ϕ*_*w*_, obtained very similar results when the multiplicity of infection (MOI) is low, around 0.35 on average (Zeng et al., 2010) – but see the **Discussion** for the effect of MOI. We estimate a 12-fold decrease for the probability of lysogenization of the virulent strain *ϕ*_*m*_ and, to our knowledge, this parameter has not been estimated elsewhere. The lysis time 1*/τ*, estimated close to 1 *h* (though with larger uncertainty), is also consistent with the biology of phage λ (De Paepe & Taddei, 2006; De Paepe et al., 2016; Lindberg et al., 2014; Shao & Wang, 2008; Zong et al., 2010). On the other hand, the adsorption rate *a* is poorly estimated. Although this issue did not arise with simulated data, our estimates based on experimental data are mostly stuck at the upper bound (10^−6^), several orders of magnitude higher than expected (De Paepe & Taddei, 2006; De Paepe et al., 2016; Lindberg et al., 2014; Shao & Wang, 2008). To investigate the sensitivity of the other estimates to the estimated value of *a*, we reiterate non-linear optimizations fixing *a* to different values ranging from 10^−9^ to 1 to compute new point estimates. The inference of most parameters was robust to the value of *a*, with no relative variation higher than 50% found for values of *a* between 10^−8^ to 1 (**Fig. S12-A**). We carry out the same sensitivity analysis for the intrinsic growth rate of the host *r* which was also poorly estimated (its distribution being mainly flat): with values of *r* ranging between 1 and 5, the probabilities of lysogenization are little affected by the perturbations in *r* (relative variation contained between −7% and +15%) while parameters *b* and *τ* – the most correlated with *r* – are the most sensitive (**Fig. S12-B**).

## 4 Discussion

In a previous study, Berngruber et al., 2013 carried out an evolution experiment with the phage λ to validate several theoretical predictions on the evolution of virulence and transmission. This study used a two-step approach: (i) a mathematical model tailored to the biology of phage λ was used to show how epidemiological dynamics feed back on the evolution of the pathogen; (ii) tracking the variation of the frequency of a viral mutant across time and across compartments (infected host and free virus) confirmed the impact of the density of susceptible cells on the transient evolution of the virus at different stages of its life cycle. This experimental validation of the theory focused on the qualitative match of the experimental results with the predictions of the model and provided empirical support for evolutionary epidemiology theory. In the present study, we adopt a reverse approach where we start from the data and try to improve the theoretical model developed to describe the evolutionary epidemiology of phage λ. We calculate the selection gradients and the differentiation across the life stages of the virus at the onset and at the end of the epidemic. This analysis allows us to better quantify the processes driving the evolutionary dynamics of the virus population.

First, we noticed that the previous model failed to properly capture the transient evolutionary dynamics in the infected compartment. We improved the goodness of fit with the experimental data by distinguishing two types of infected cells: lysogens (*L*) and cells committed to the lytic pathway (*Y*). Indeed, adding extra stages allowed us to observe a transitory peak in the frequency of hosts infected by the virulent phage (**Fig. S13-A**, middle panel), similar to what we observed in the experimental data. In this model, the peak is due to the frequency of the virulent phage among *Y* bacteria that, unlike *L* bacteria, transiently overshoots during the acute phase of the epidemic. Crucially, lysis was assumed to be instantaneous in the original model (Berngruber et al., 2013) and including this additional stage in the phage life cycle allowed us to take the lag between infection and lysis into account (Brown et al., 2022; De Paepe et al., 2016; Geng et al., 2023; Mitarai et al., 2016; Yin & McCaskill, 1992; You & Yin, 1999). For the sake of simplicity and parsimony, we only kept a single extra stage *Y*, yielding exponentially distributed lysis time (Mitarai et al., 2016).

Second, we performed a theoretical analysis of this new model to go beyond the numerical approach used in Berngruber et al., 2013. In particular, we obtained analytic approximations for the spread of the virulent phage at the beginning and at the end of the epidemic: (i) the virulent mutant increases in frequency at the onset of the epidemic (when the density of susceptible hosts is high) with a speed approximately proportional to −Δ*ϕ*; (ii) the virulent mutant decreases in frequency at the end of the epidemic (when the density of susceptible hosts is low) with a negative selection gradient −Δ*α*, which is consistent with the prediction that in the long-term, and in the absence of an influx of susceptible hosts, the temperate phage should evolve a fully lysogenic strategy where *α* = 0 and *ϕ* = 1 (Bruce et al., 2021; Wahl et al., 2019); (iii) the virulent phage is always more frequent in the the free virus stage than among prophages and this differentiation is approximated by 1 + Δ*α/α*_*w*_ at the end of the epidemic. The first approximation only held for a very limited period of time because the density of susceptible cells drops very rapidly and this epidemiological feedback affects the selection on transmission and virulence. In contrast, the final two predictions were valid as soon as the epidemic reaches high prevalence and, interestingly, yielded a novel way to estimate the rates of prophage reactivations of both strains.

Third, we developed a statistical inference approach to estimate the parameters of the model. We implemented a two-step MLE procedure: (i) we first obtained point estimates of the reactivation rates *α*_*w*_ and *α*_*m*_ of both viral strains, (ii) then we fixed *α*_*w*_ and *α*_*m*_ to their point estimates and ran non-linear optimizations to infer the remaining parameters of the model (except the burst size *B* which had to be fixed because of an identifiability issue). The Sieve bootstrap technique was used to generate joint distributions for all the estimated parameters (**Table 2**). We showed that our estimates of the key phenotypic traits *α*_*w*_, *α*_*m*_ and *ϕ*_*w*_ were consistent with existing literature (**Table S3**).

Crucially, our inference approach is very different from previous studies. First, we analyse the dynamics of a polymorphic viral population. Second, we use three different types of data to infer the parameter values of the model: (i) epidemiological data (i.e. the prevalence of the infection), (ii) temporal variation of variant frequencies and (iii) differentiation of the variant frequency across compartments. Each data type carries complementary information; together, they allow us to jointly estimate the life-history traits of both strains of the phage. This novel method is particularly well suited to estimate the rates of prophage reactivation, for which only the endemic treatment is needed – see **SI Appendix §S3.2** where we propose a Bayesian counterpart to easily obtain posterior distributions and credible intervals for these two parameters (applied to our experimental data, we show results in **Fig. S14-A**). Estimating the probabilities of lysogenization is however more difficult as it requires us to monitor the transient dynamics of the epidemic treatment. The rapid epidemiological feedback that occurred in emerging epidemics makes it necessary to consider both the epidemiology and the evolution of the infection. It might thus be relevant to carry out shorter experiments for the epidemic treatment – i.e., focusing on the the early state of the epidemic – but with a more frequent sampling effort to monitor the change in frequency more precisely. Alternatively, the use of two-stage chemostats, where the influx of susceptible hosts could be maintained experimentally (Bonachela & Levin, 2014; Husimi et al., 1982), might also be an option worth investigating.

Even though we have improved the original model, many features of the present model could be challenged. For example, we assume all phenotypic traits to be constant across time and across chemostats. Yet, key life-history traits of phage λ are known to vary with the environment (Wegrzyn & Wegrzyn, 2005) (i.e., phenotypic plasticity). In particular, the probability of lysogenization and the reactivation rate of the λ mutant used in the evolution experiment is temperature-sensitive (Berngruber et al., 2013; St-Pierre & Endy, 2008; Zong et al., 2010). Interestingly, the estimated rates of reactivation varied among chemostats (**Fig. S14-B**, using a Bayesian approach). This variation may have been driven by small variations in temperature among the chemostats that could have affected the reactivation rate of the mutant (higher rates of reactivation would be expected with higher temperature). The probability of lysogenization of phage λ is also known to be more likely to occur at high multiplicity of infection (MOI) (Kourilsky & Knapp, 1975; Kourilsky, 1973; Zeng et al., 2010). MOI-dependent phenotypic plasticity may also affect the rate of reactivation of temperate phages (Bruce et al., 2021), or the lysis time of virulent phages (Rutberg & Rutberg, 1965). Yet, our model has the advantage of the parsimony, while already convincingly reproducing the qualitative patterns observed in the data. More sophisticated models would improve the fit to the data compared to simpler, nested models. Additional details may be rooted in experimental knowledge on the system, but the data may not be rich enough to infer the extra-parameters precisely (and accordingly, these more complex models may not be selected over simpler ones in statistical model comparisons). Moreover, simpler models are easier to analyze mathematically and lend themselves more easily to interpretation. It is thus delicate to know where to draw the line between the parsimony and the goodness of fit of a model. Navigating in this trade-off, a first model was developed capturing the transient increase in virulent strain (Berngruber et al., 2013), and we extended it with the known delay between phage infection and lysis to better capture the similarity in variant frequency in cells and free viruses. Our final model fits the data well, but not all parameters can be inferred precisely, in ways that depend on the details of the model and would have been difficult to anticipate before formally fitting the model. Our study emphasizes the benefits of combining theoretical, statistical and experimental approaches. Combining these perspectives effectively requires an iterative process. Each step is an opportunity to challenge and improve our understanding of the biological model, to elucidate microbial life cycles and to provide support and guidance during experimentation. While most experimental life-history assays only focus on monomorphic population, we demonstrate the relevance of evolution experiments where different strains are put in competition and where tracking their relative frequencies over time may yield novel and efficient ways to estimate some dimensions of their phenotypes, especially when it is difficult to access absolute densities. This approach could be used under different conditions and in particular *in vivo* (De Paepe et al., 2016). The present study may thus help characterize the phenotypic traits of microbial organisms used in experimental evolution. More broadly, this overall quantitative process is similar in spirit with what can be done in epidemiological studies, particularly in public health – though generally with a much simpler description of the life cycle. During the COVID-19 pandemic, for instance, the successive emergence and sweeps of SARS-CoV-2 variants of concern has raised many questions about the short and long-term evolution of the virus, especially in terms of transmission and virulence. Statistical inference based on demographic/epidemiological data (e.g., prevalence, deaths, hospital admissions) and genetic data (e.g., strain frequencies) quantified variants properties such as transmission, virulence, infectious period, immune escape or generation interval distribution – e.g., (Benhamou et al., 2023; Blanquart et al., 2022; Davies et al., 2021). In many scenarios, the frequency of a variant could be measured in different compartments, for example between naive and vaccinated hosts (e.g., for SARS-CoV-2 variants Omicron), or between infected hosts and an environmental compartment (e.g. enteropathogenic *E. coli* in environmental waters). As shown in this study, interpreting the temporal variation in the selection gradient and the differentiation of a novel variant between distinct compartments may be a powerful tool to clarify the relationship between its frequencies in distinct life stages and to better characterize its phenotype.

## Abbreviations (alphabetical order)

AIC: Akaike Information Criterion
ARMA: AutoRegressive-Moving-Average
CFP: Cyan Fluorescent Protein
CI: Confidence Interval
COVID-19: Coronavirus Disease 2019
FACS: Fluorescence-Activated Cell Sorting
i.i.d.: independent and identically distributed
MLE: Maximum Likelihood Estimation
MOI: Multiplicity Of Infection
ODE: Ordinary Differential Equation
qPCR: quantitative Polymerase Chain Reaction
SARS-CoV-2: Severe Acute Respiratory Syndrome Coronavirus 2
SD: standard deviation
YFP: Yellow Fluorescent Protein

## Supplementary material

**Supplementary Figures and Tables** (PDF)

**Supplementary Information (SI Appendix)** (PDF)

## Data accessibility

Data that were used in this study, along with the scripts for the analyses (in R), are available in this GitHub repository.

## Funding

We acknowledge the French Ministry of Higher Education and Research and the École Normale Supérieure Paris-Saclay for the PhD scholarship of WB.

## Conflict of interest

The authors declare no conflict of interest.

## Notes

### Competing Interest Statement

The authors have declared no competing interest.

### Summary of Updates

This version of the manuscript has been revised to update the main text.

