## Supplementary figures and tables for "Evolution of Virulence in Emerging Epidemics: From Theory to Experimental Evolution and Back"

July 26, 2024

**Supplementary figures** (pp. 2-15)

– **Figure S1:** Numerical simulations for the epidemic and endemic treatments.
– **Figure S2:** Simulated datasets differing in quantity and/or quality.
– **Figure S3:** Approximation of the selection gradient of the virulent phage at the early stage of the epidemic.
– **Figure S4:** Differentiation of the virulent phage.
– **Figure S5:** Point estimates of the reactivation rates $\alpha_w$ and $\alpha_m$ from simulated datasets.
– **Figure S6:** Log-likelihood landscape according to the values of parameters $b$ and $B$.
– **Figure S7:** Parameter point estimates from simulated data.
– **Figure S8:** Density distributions of parameter estimates.
– **Figure S9:** Pairwise correlations.
– **Figure S10:** Fitted values from sieve bootstrap on experimental data.
– **Figure S11:** Sensitivity of the inference of estimated parameters to the fixed burst size.
– **Figure S12:** Sensitivity of the estimated parameters to perturbations in the adsorption rate or in the bacterial intrinsic growth rate.
– **Figure S13:** Comparisons between different lysis time distributions.
– **Figure S14:** Bayesian inference of the rates of prophage reactivations.

**Supplementary tables** (pp. 16-17)

– **Table S1:** Parameter values used in the simulations.
– **Table S2:** Bounds for non-linear optimizations.
– **Table S3:** Examples of values for phage parameters from previous studies.

**Supplementary references** (p. 18)

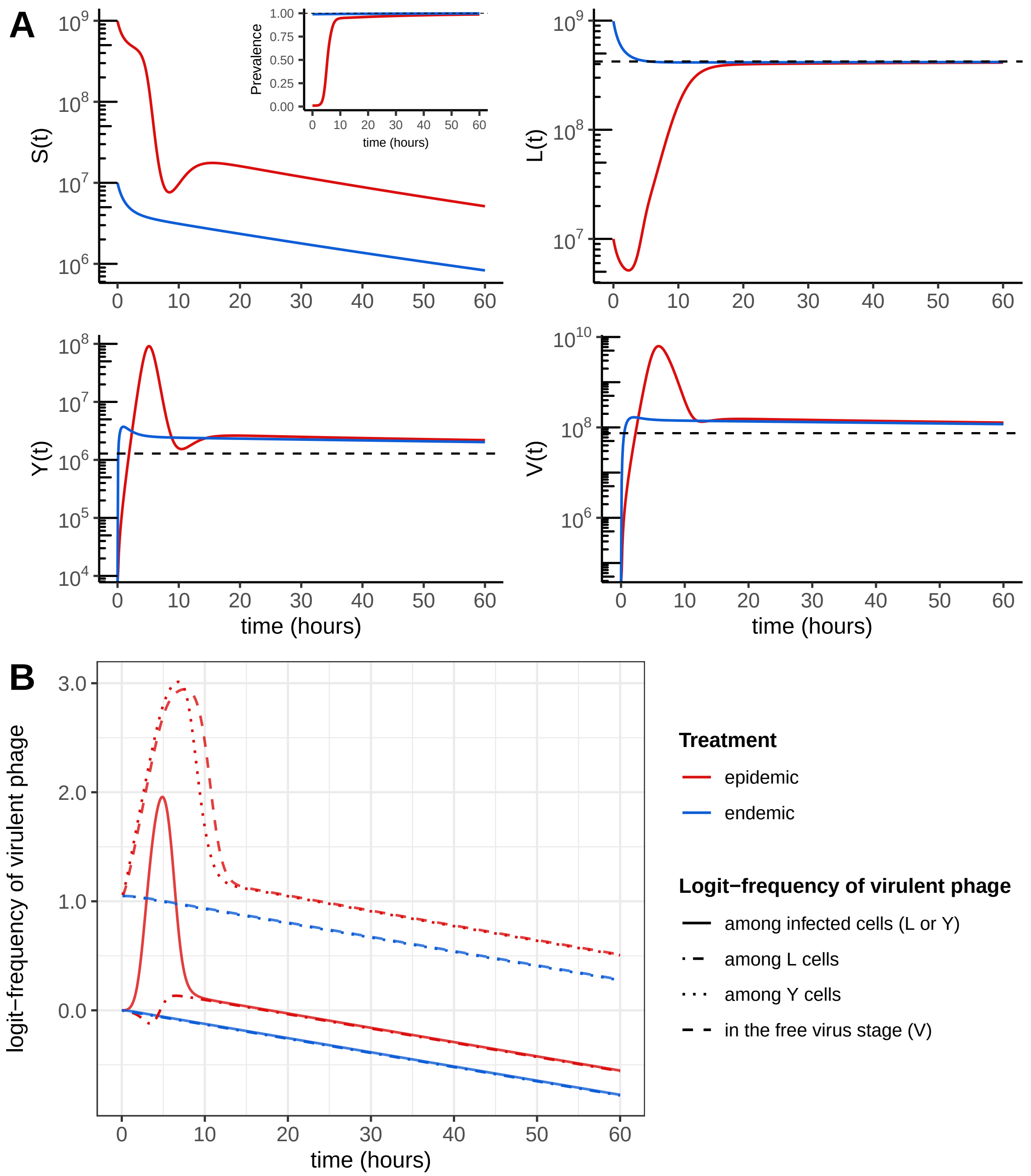


Figure S1: **Numerical simulations for the epidemic and endemic treatments.** (A) Total density of each compartment (horizontal dashed lines indicate endemic equilibrium values) and (B) logit-frequencies of the virulent strain $m$. See **Table S1** for parameter values. At $t = 0$, bacteria are at carrying capacity $K$ with initial prevalence 1% (epidemic treatment) or 99% (endemic treatment). The initial prophage ratio for the two strains is 1:1.

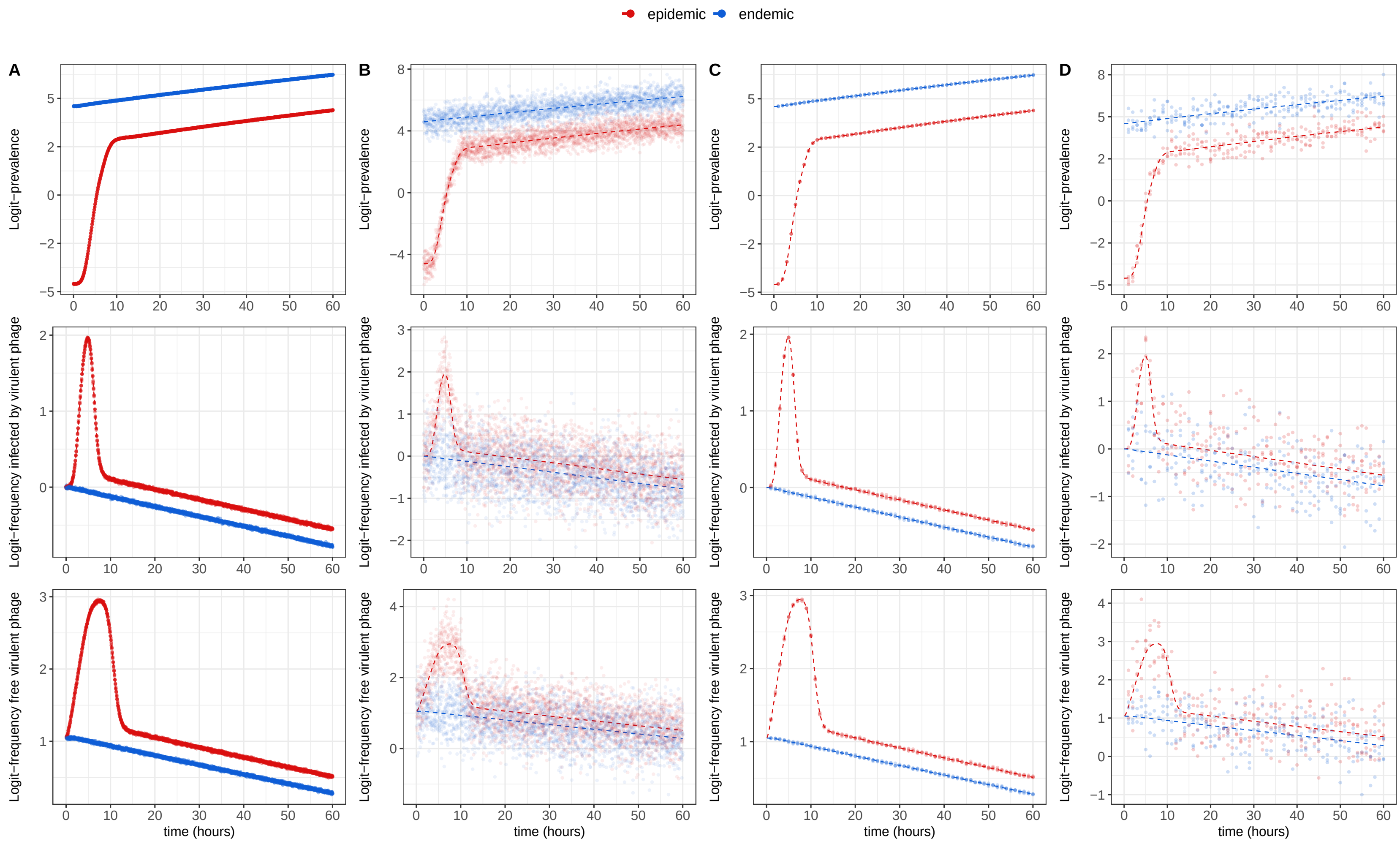


Figure S2: **Simulated datasets differing in quantity and/or quality.** These four sets of simulated data (points) stem from the same two deterministic simulations (epidemic vs. endemic treatment, in dashed lines, see **Table S1** for parameter values and initial conditions) to which we add white Gaussian noise to mimic measurement errors. This was performed four times for each simulation to generate four replicates (chemostats) per treatment. Simulated datasets differ in terms of sampling effort: 1 (C-D) vs. 10 $h^{-1}$ (A-B), and/or accuracy: SD of measurement errors = 0.01 (A-C) vs. 0.5 (B-D).

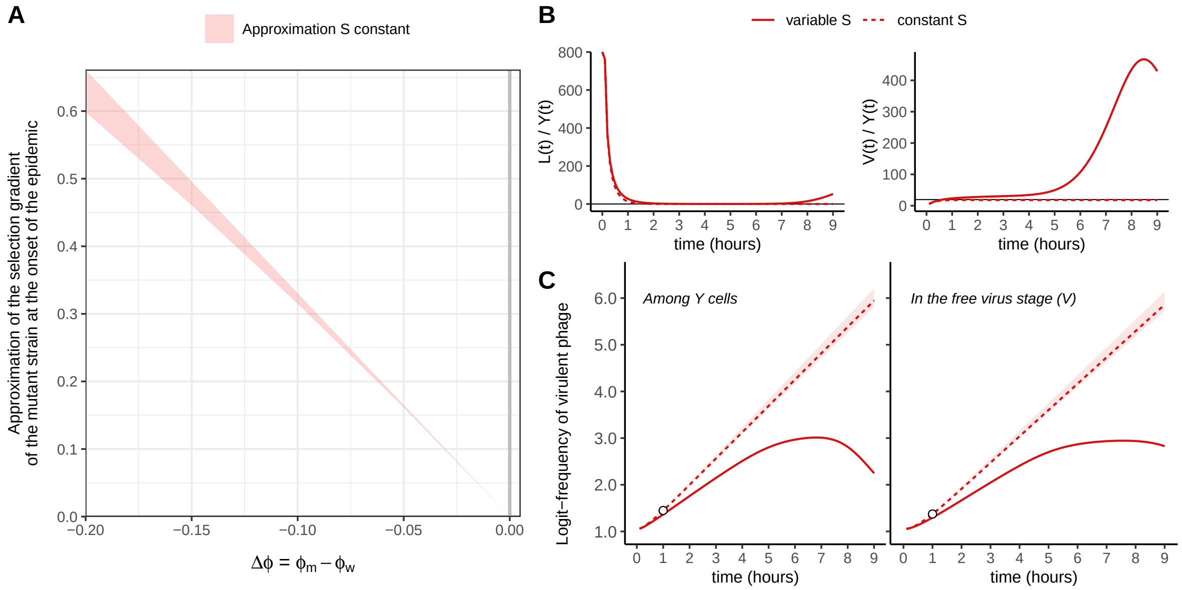


Figure S3: **Approximation of the selection gradient of the virulent phage at the early stage of the epidemic.** At $t = 0$, bacteria are at carrying capacity $K$ with initial prevalence 1% (epidemic treatment) and initial prophage ratio for the two strains 1:1. We investigate the case where the pool of susceptible host is kept constant to its initial value $S(t = 0) = 0.99 \times K$ throughout the course of infection (dashed lines). See **Table S1** for parameter values. In panel A, we vary $\Delta\phi = \phi_m - \phi_w$ by varying $\phi_m$; in panels B and C, $\Delta\phi = -0.18$. (A) Approximation interval for the selection gradient of the mutant strain (slope of logit($f(t)$)) against $\Delta\phi$, as given in **SI Appendix §S2.2** (equation (S17)); note that the selection gradient is positive when $\Delta\phi$ is negative, meaning that the virulent phage is predicted to be selected for at the early stage of the epidemic. (B) Temporal dynamics of the ratios $L(t)/Y(t)$ and $V(t)/Y(t)$ along with their approximations (horizontal black lines). (C) Predictions of the trajectories of the logit-frequencies of the virulent phage among lytic cells ($Y$) and in the free virus stage ($V$), starting from values at $t = 1$ (white points). Indeed, at $t = 0$, only lysogens are introduced in the susceptible population, so we first let the phage-bacteria system reaches its new dynamical regime – when epidemiological dynamics have reached (quasi-)equilibrium in panel B. The shaded area derives from the approximation interval illustrated in panel A

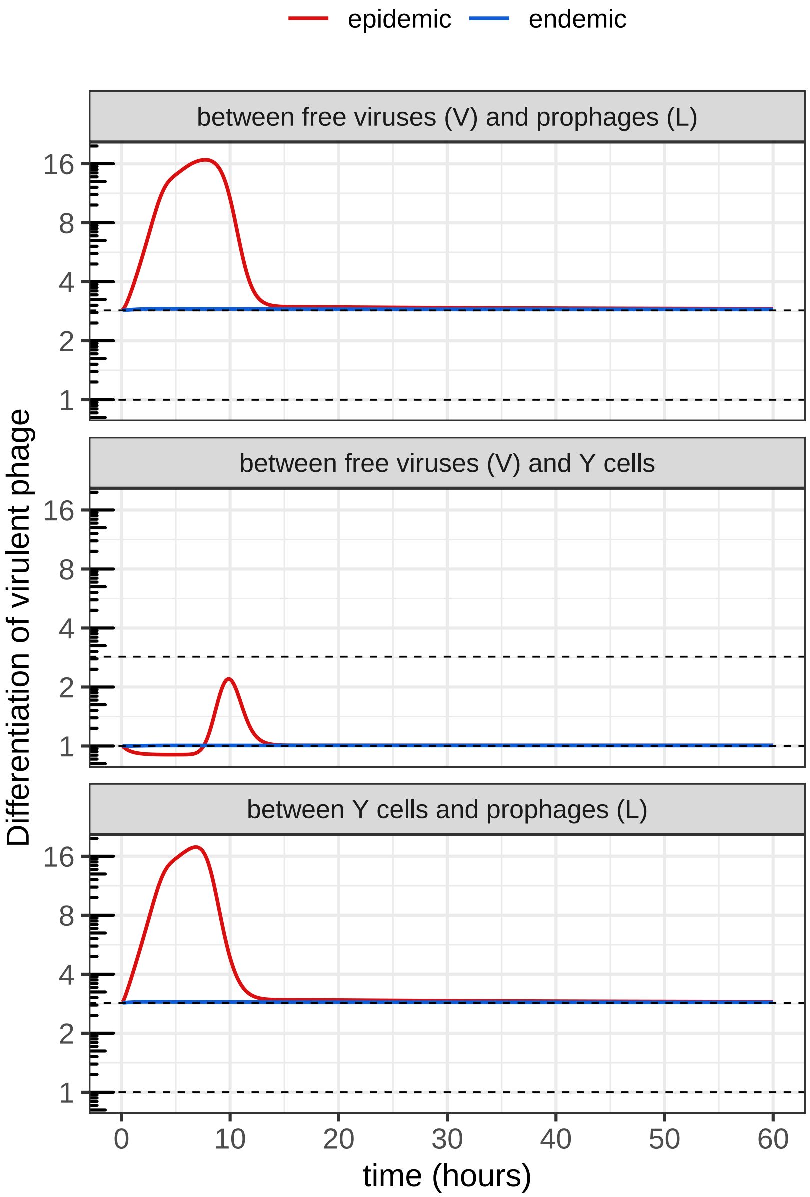


Figure S4: **Differentiation across distinct life stages of the virus.** In **SI Appendix §S2.4**, we show that the differentiation between free viruses ($V$) or lytic cells ($Y$) and prophages ($L$) is predicted to converge towards approximately $1 + \Delta\alpha/\alpha_w > 1$ (uppermost dashed line), and that the differentiation between free viruses ($V$) and lytic cells ($Y$) towards approximately 1 (no differentiation, lowermost dashed line). In the endemic case – including the late stage of the epidemic –, the virulent strain is therefore more frequent among free viruses and lytic cells than among prophages but is as frequent among free viruses as among lytic cells. See **Table S1** for parameter values and initial conditions.

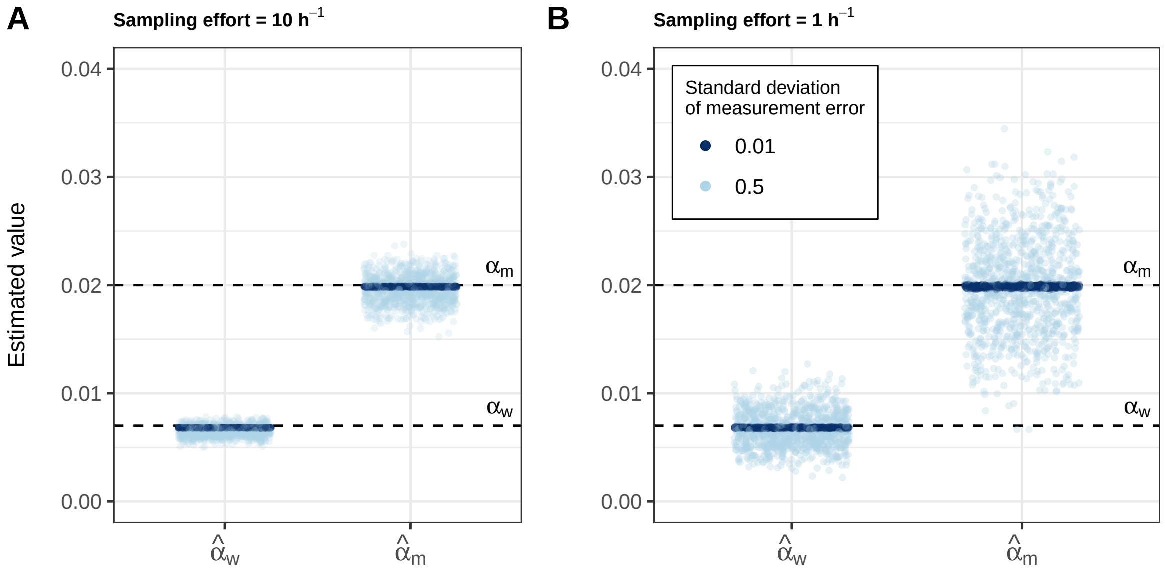


Figure S5: **Point estimates of the reactivation rates $\alpha_w$ and $\alpha_m$ from simulated datasets.** We consider data with different sampling efforts ((A) 10 vs. (B) 1 $h^{-1}$) and/or SD of measurement errors (0.01 (dark blue) vs. 0.5 (light blue)); for each combination, we computed 1,000 simulated datasets by adding Gaussian noise to the same two simulations (epidemic vs. endemic) shown in **Fig. S1**; repeating this four times, we independently generate four replicates (chemostats) per treatment. For each replicate, we only keep the data from the time point the system had reached a prevalence of 95%. From each simulated dataset, we eventually compute point estimates of both parameters $\alpha_w$ and $\alpha_m$ (see **Materials and methods §2.3.1** and **SI Appendix §S3.1**) which overall show a good match with the values used in the original simulation (horizontal dashed lines).

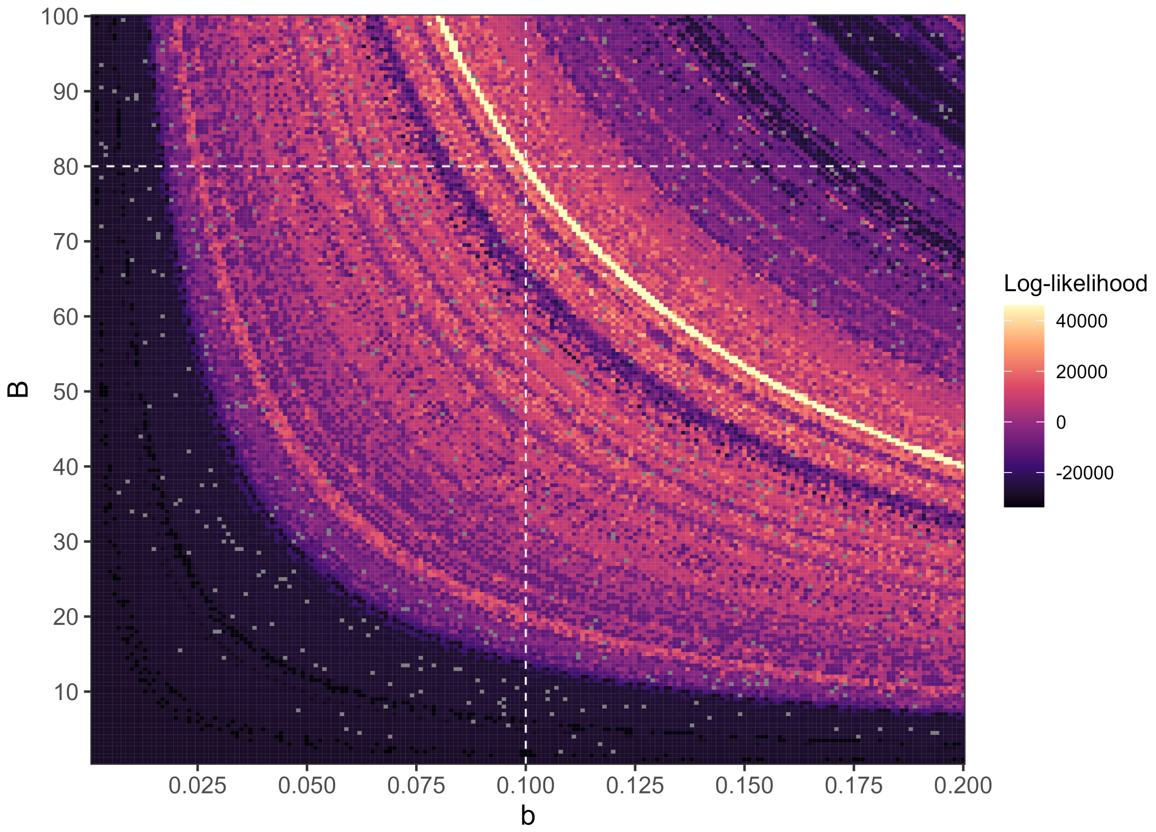


Figure S6: **Log-likelihood landscape according to the values of parameters $b$ and $B$.** We used the simulated dataset closest to the original deterministic simulation (sampling effort = 10 $h^{-1}$ and SD of measurement error = 0.01, see **Fig. S2-A**). For each pair $(b,\ B) \in [0,\ 0.2] \times [0,\ 100]$, we also fixed the rates of prophage reactivation $\alpha_w$ and $\alpha_m$ (as though correctly estimated beforehand) as well as $K$ and $\delta$ while we maximized the overall log-likelihood over the remaining parameters. White dashed lines indicate the values used in the original simulations. Unsuccessful completions are shown in grey. This landscape points out that parameters $b$ and $B$ are not separately identifiable; its shape suggests that only the product $b \times B$ is identifiable, in particular the log-likelihood always reaches the highest value (lightest curve) when $b \times B$ is around 8.

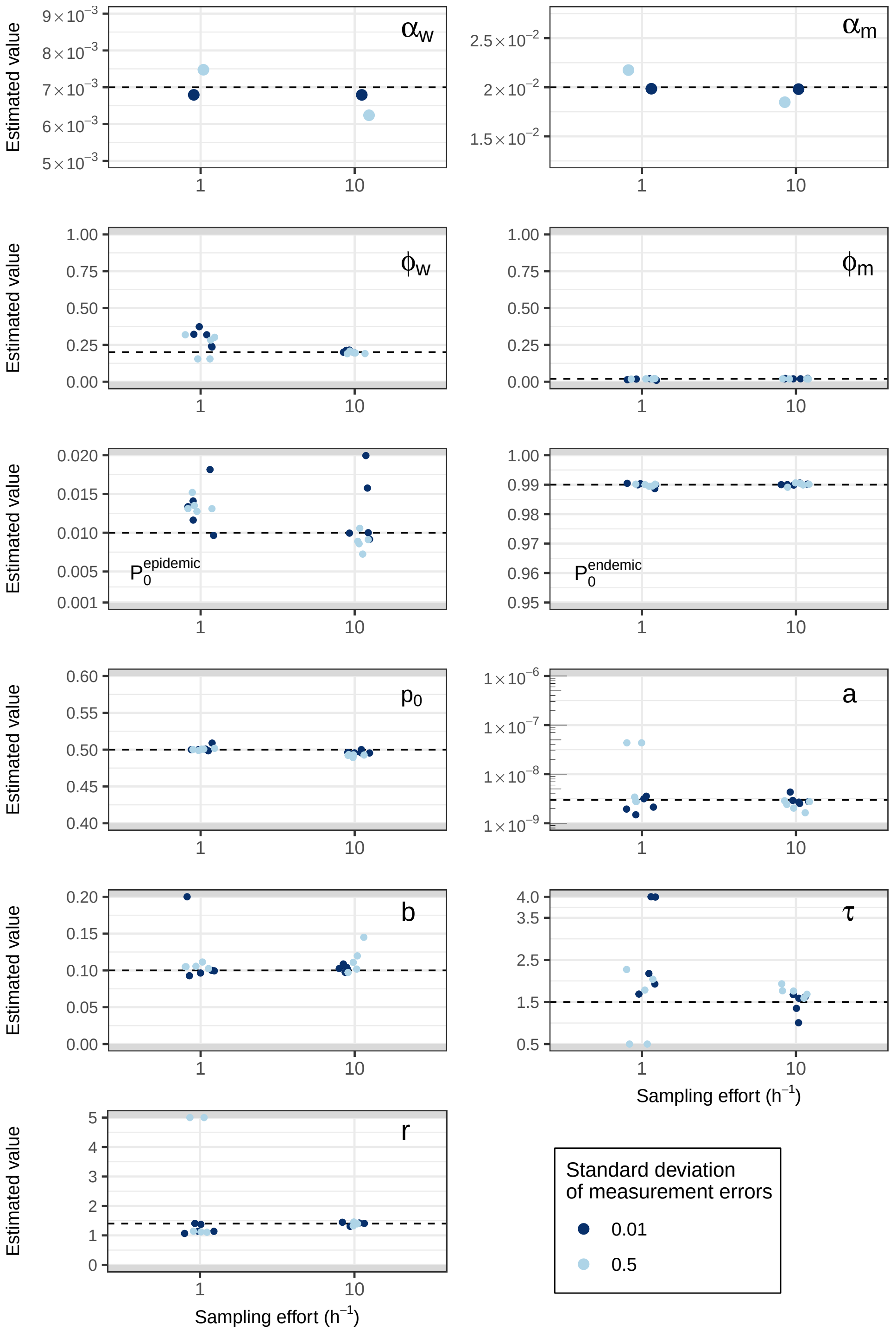


Figure S7: **Parameter point estimates from simulated data.** We use the four simulated datasets described in **Fig. S2**, which all stem from the same deterministic simulation but differ in sampling effort (1 vs. 10 $h^{-1}$) and/or SD of measurement errors (0.01 vs. 0.5). For each simulated dataset, we first estimate $\alpha_w$ and $\alpha_m$; we then run non-linear optimizations to estimate the remaining parameters ($K$, $\delta$ and $B$ are fixed), starting from a set of 2,000 initial values and keeping parameter values from the best fit. We repeat this procedure for five sets of initial values, yielding five point estimates for each remaining parameter. Dashed horizontal lines refers to the values we use to generate simulated data and grey areas indicate out-of-bounds ranges of values.

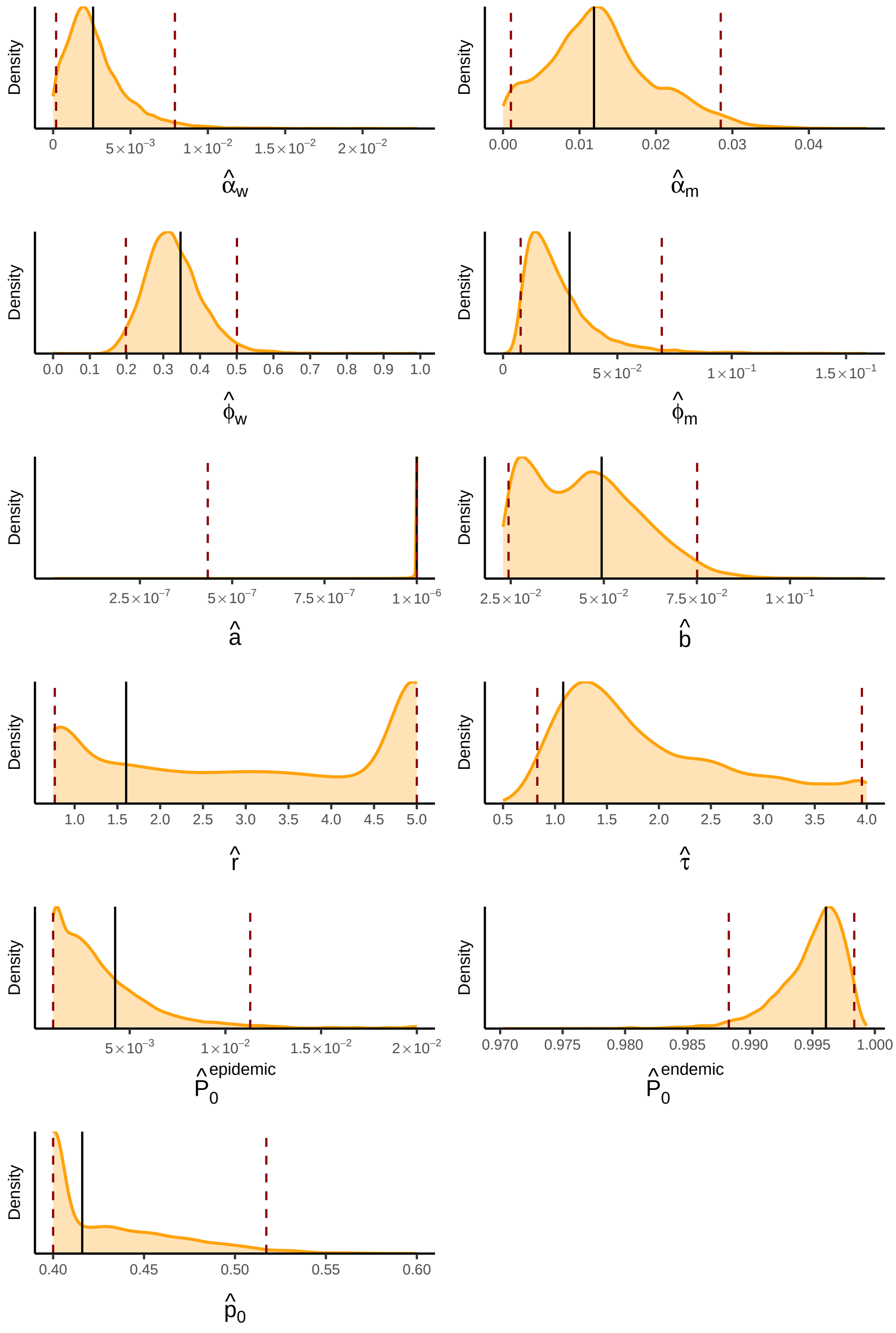


Figure S8: **Density distributions of parameter estimates.** Using sieve bootstrap on the residuals between experimental data and the best fit of our model (black vertical solid lines), we generate $1,000$ new datasets on which we reiterate the estimation procedure to compute the joint distributions of estimated parameters; we only keep results with successful convergence ($n = 6,686$). Dashed vertical lines indicates 2.5% and 97.5% quantiles.

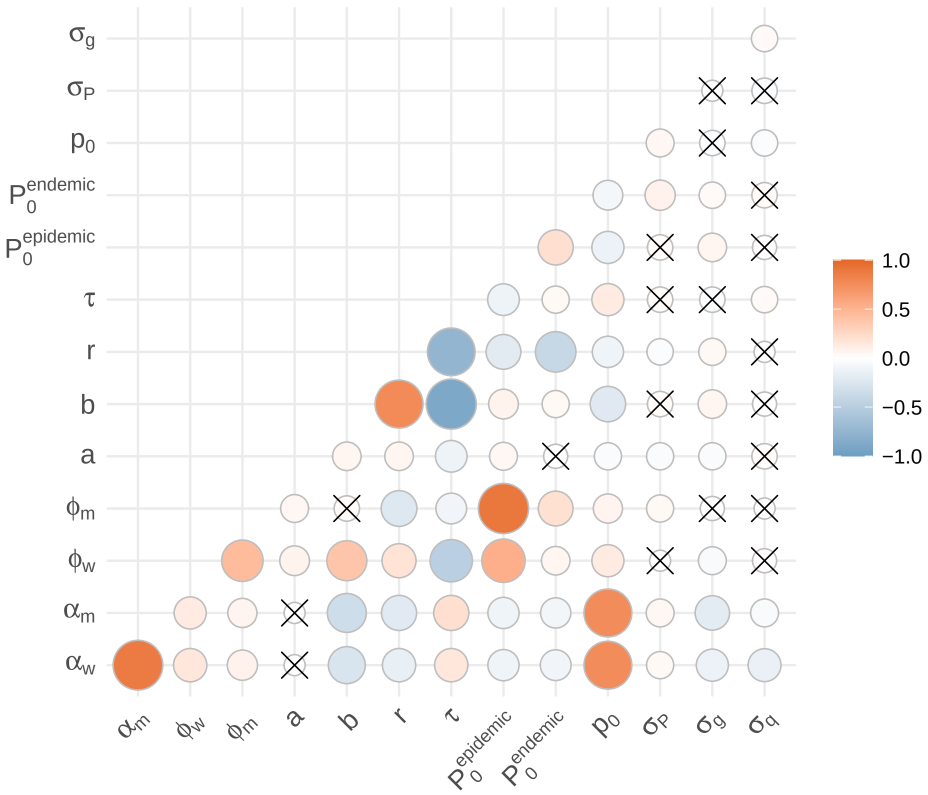


Figure S9: **Pairwise correlations.** We compute Pearson correlation coefficient of bootstrap-based distributions of parameter estimations (cf. **Fig. S8**, $n = 6,686$). Crosses indicate non-significant coefficients ($p$-value $> 0.05$).

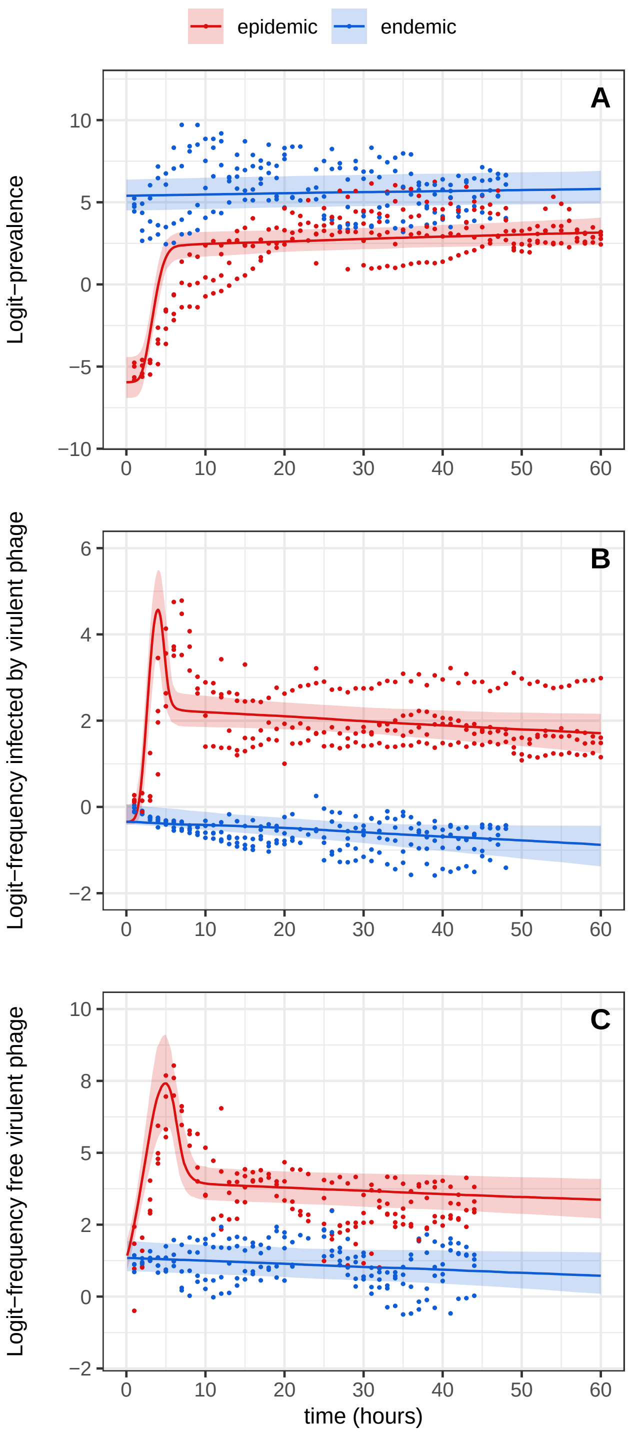


Figure S10: **Fitted values from sieve bootstrap on experimental data.** Distributions of fitted values – median (line) and 95% interval (shaded envelope) – were obtained using sieve bootstrap on the residuals between experimental data (points) and the best fit of our model. Initial prevalence is either low (endemic condition, in blue) or high (epidemic condition, in red). (A) Logit-prevalence of infection; (B) logit-frequency of cells infected (either lysogenic or lytic) by the the mutant (virulent) phage; (C) logit-frequency of the mutant (virulent) in the culture medium (free virus stage).

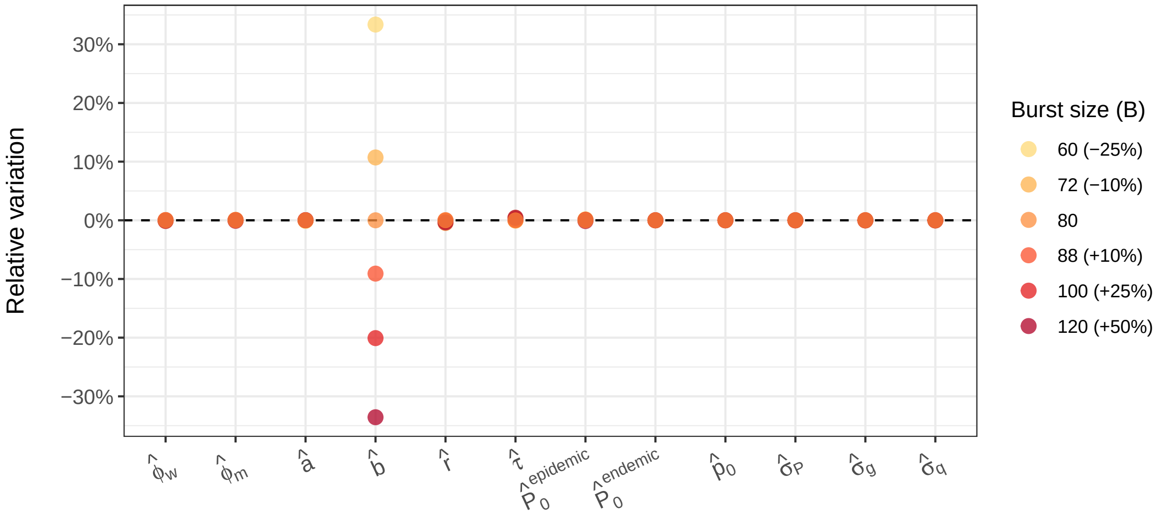


Figure S11: **Sensitivity of the inference of estimated parameters to the fixed burst size.** We apply ±10%, ±25% and +50% perturbations on the fixed burst size $B$ (original fixed value: $B = 80\ virus\,.\,cell^{-1}$) and we reiterate non-linear optimizations as before to compute new point estimates. Relative variations (y-axis) refer to the percentage of variation of these new estimates compared to the original best MLE estimates (without perturbation). All parameters show high robustness to these perturbations, except for parameter $b$ as expected from **Fig. S6** (but the product $b \times B$ is always around 3.95).

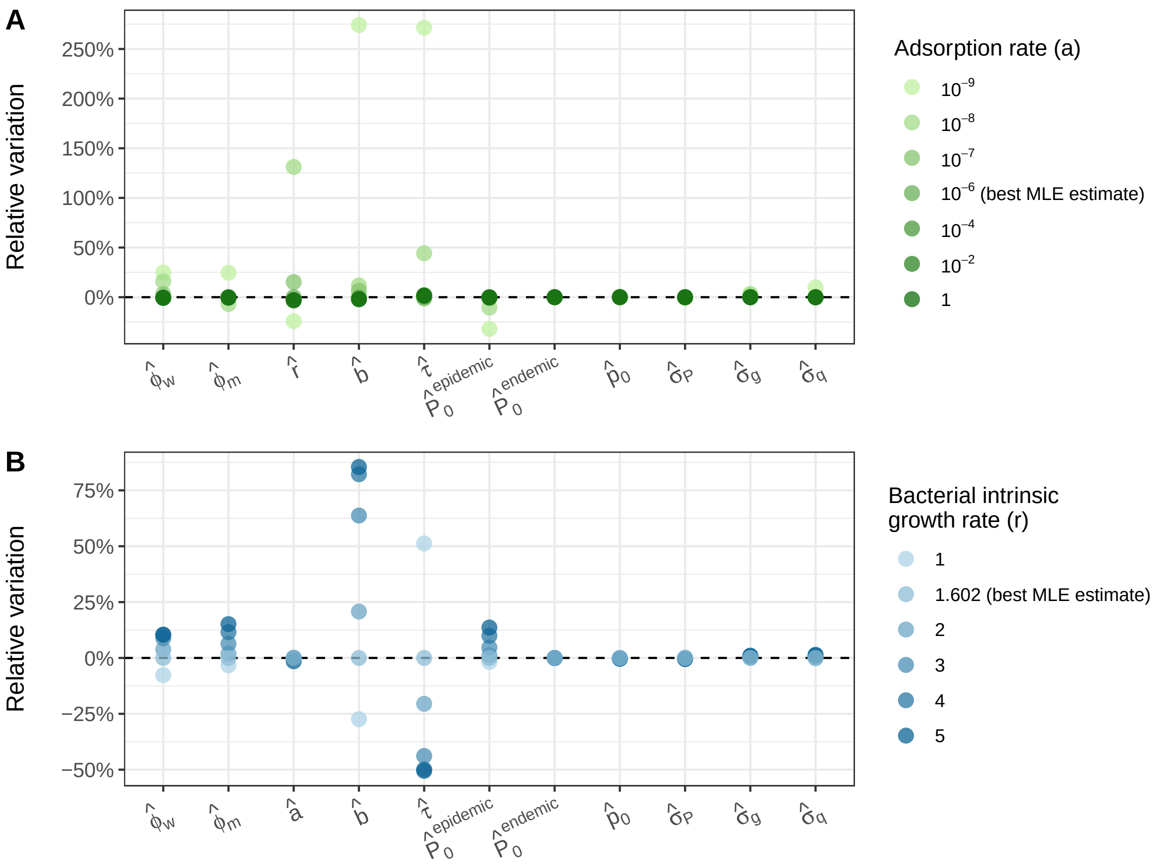


Figure S12: **Sensitivity of the estimated parameters to perturbations in the adsorption rate or in the bacterial intrinsic growth rate.** Both these parameters – $a$ and $r$, respectively – were poorly estimated with experimental data. We thus investigate *a posteriori* how the other estimated parameters are impacted when $a$ and $r$ deviate from their best MLE estimates (both expressed in $h^{-1}$). Iteratively fixing $a$ (A) or $r$ (B) to some values, we then reiterate non-linear optimizations as before to compute new point estimates. Relative variations (y-axis) refer to the percentage of variation of these new estimates compared to best MLE estimates.

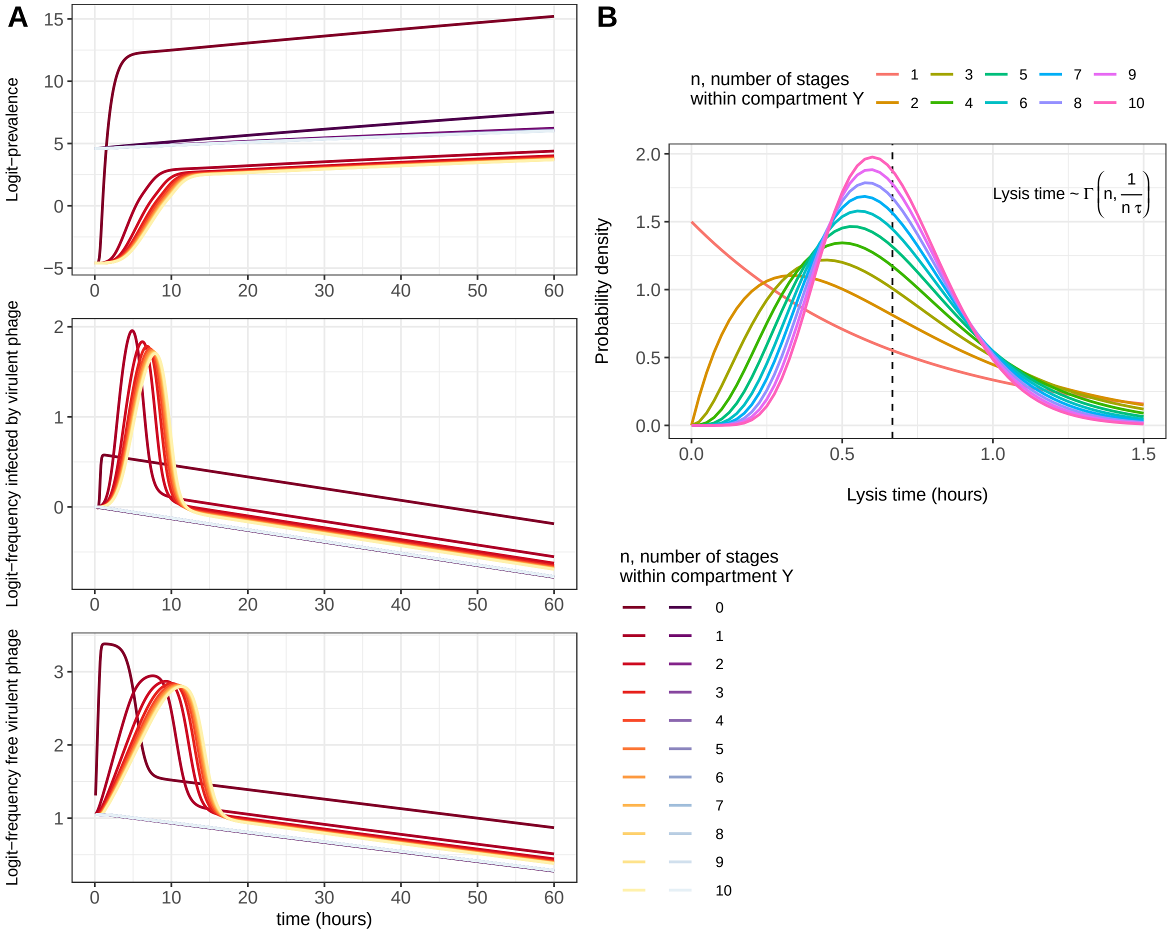


Figure S13: **Comparisons between different lysis time distributions.** Using linear/Gamma chain trick, compartments $Y$ (both $Y_w$ and $Y_m$) are stratified into $n$ successive stages. As ODEs implicitly assume exponentially distributed sojourn time, the lysis time is thus the sum of $n$ i.i.d. exponential distributions, that is a gamma distribution with shape parameter $n$ and scale parameter $1/(n\tau)$. The case $n = 1$ corresponds to the exponential distribution – which is the one we use in the main text – and we also include the case $n = 0$ (no compartment $Y$), as in Berngruber et al., 2013. See **Table S1** for parameter values. The mean lysis time is given by $1/\tau$ (vertical dashed line in B). At $t = 0$, bacteria are at carrying capacity $K$ with initial prevalence 1% (epidemic treatment) or 99% (endemic treatment). The initial prophage ratio for the two strains is 1:1.

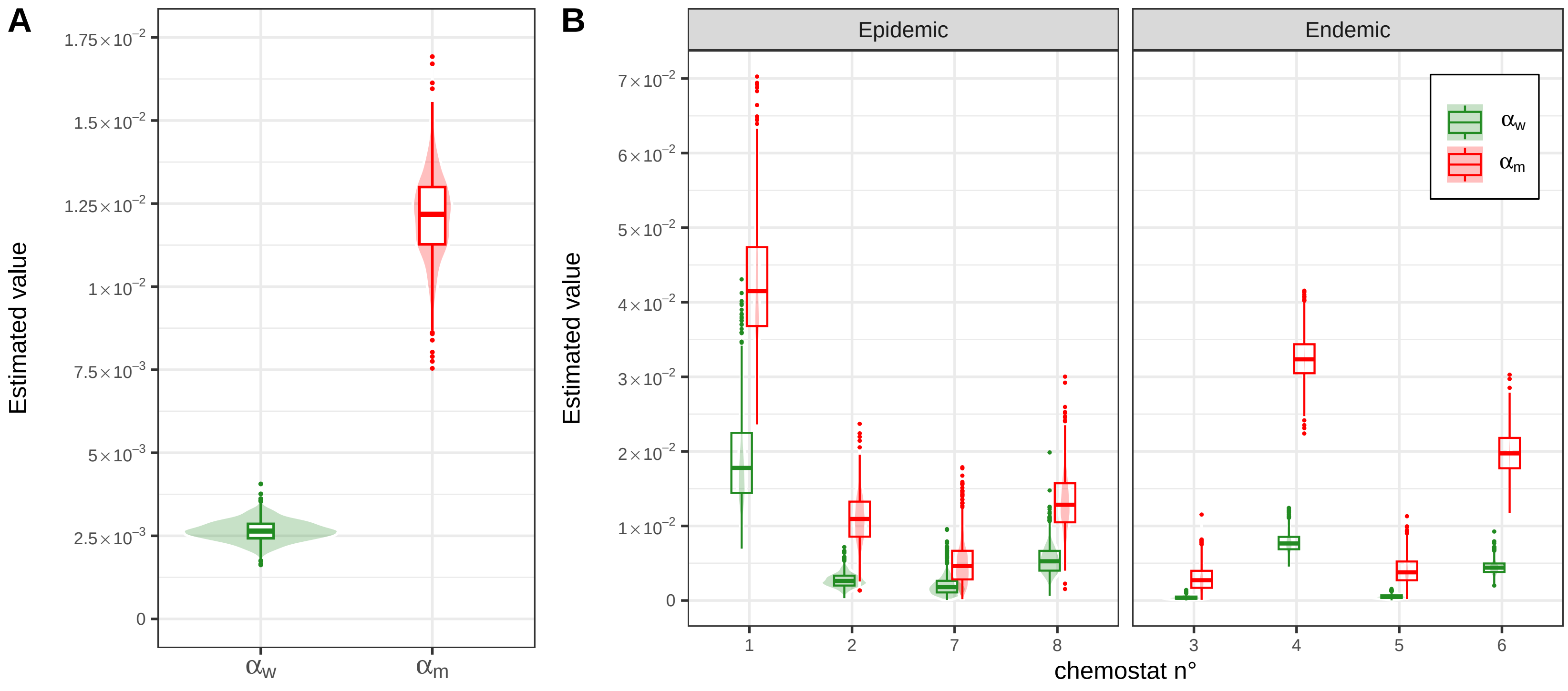


Figure S14: **Bayesian inference of the rates of prophage reactivations.** We estimate $\alpha_w$ and $\alpha_m$ from experimental data using a Bayesian approach (see details in **SI Appendix §3.2**). (A) Parameters $\alpha_w$ and $\alpha_m$ are assumed to be the same for all chemostats – $\hat{\alpha}_w = 2.65 \times 10^{-3}$ (mean, 95% credible interval $[2.03 \times 10^{-3},\ 3.32 \times 10^{-3}]$) and $\hat{\alpha}_m = 1.21 \times 10^{-2}$ (mean, 95% credible interval $[9.48 \times 10^{-3},\ 1.47 \times 10^{-2}]$) –; (B) parameters $\alpha_w$ and $\alpha_m$ are allowed to vary across chemostats.

Table S1: **Parameter values used in the simulations.** The subscripts $w$ and $m$ refer to the wildtype strain and the mutant (or virulent) strain $\lambda$cI857 of phage $\lambda$, respectively. $P_0^{epidemic}$ and $P_0^{endemic}$ correspond to the initial conditions (at $t = 0$) of the prevalence for the endemic and epidemic treatment, respectively; $p_0$ corresponds to the initial condition of the frequency of lysogenic hosts ($L$) infected by the virulent phage (infected cells are all lysogenic at $t = 0$). See **Table 1** for notations.

| **Parameter** | **Value** | **Unit** |
|---|---|---|
| $P_0^{epidemic}$ | 1% | – |
| $P_0^{endemic}$ | 99% | – |
| $p_0$ | 0.5 | – |
| $\alpha_w$ | $7 \times 10^{-3}$ | $h^{-1}$ |
| $\alpha_m$ | $2 \times 10^{-2}$ | $h^{-1}$ |
| $\phi_w$ | 0.2 | – |
| $\phi_m$ | $2 \times 10^{-2}$ | – |
| $a$ | $3 \times 10^{-9}$ | $h^{-1} \,.\, cell^{-1}$ |
| $b$ | 0.1 | – |
| $\tau$ | 1.5 | $h^{-1}$ |
| $B$ | 80 | $virus \,.\, cell^{-1}$ |
| $r$ | 1.4 | $h^{-1}$ |
| $K$ | $10^9$ | $cell$ |
| $\delta$ | 0.8 | $h^{-1}$ |
| $\sigma_P,\ \sigma_g,\ \sigma_q$ | 0.01/0.5 | – |

Table S2: **Bounds for non-linear optimizations.** Bounds are placed on estimated parameters to constrain optimizations to relevant ranges of values. Besides, starting values are uniformly drawn between these bounds.

| **Parameter** | **Lower bound** | **Upper bound** | **Unit** |
|---|---|---|---|
| $P_0^{epidemic}$ | 0.95 | 1 | – |
| $P_0^{endemic}$ | $10^{-3}$ | $2 \times 10^{-2}$ | – |
| $p_0$ | 0.4 | 0.6 | – |
| $\phi_w,\ \phi_m$ | 0 | 1 | – |
| $a$ | $10^{-9}$ | $10^{-6}$ | $h^{-1} \,.\, cell^{-1}$ |
| $b$ | 0 | 0.2 | – |
| $r$ | 0 | 5 | $h^{-1}$ |
| $\tau$ | 0.5 | 4 | $h^{-1}$ |
| $\sigma_P,\ \sigma_g,\ \sigma_q$ | $10^{-3}$ | 2 | – |

Table S3: **Examples of values for phage parameters from previous studies.** We do not include the probability of fusion $b$, as it is not separately identifiable from the burst size $B$ in our model and as it is not really considered in most studies. We also do not include the probability of lysogenization of the virulent phage $\phi_m$ as, to our knowledge, it has not been estimated elsewhere.

| **Parameter** | **Value** | **Unit** | **Reference** |
|---|---|---|---|
| $\alpha_w$ | $3.4 \times 10^{-4}$ | $h^{-1}$ | (De Paepe et al., 2016) |
| | $6.3 \times 10^{-3} - 3.6 \times 10^{-2}$ | | (De Paepe et al., 2016) |
| | $2.4 \times 10^{-5}$ | | (Little et al., 1999) |
| | $\sim 10^{-7} - 10^{-6}$ | | (Zong et al., 2010) |
| $\alpha_m$ | $1.3 \times 10^{-2}$ $^{(i)}$ | $h^{-1}$ | (De Paepe et al., 2016) |
| | $\sim 10^{-3}$ | | (Zong et al., 2010) |
| $\phi_w$ | 0.19 | – | (De Paepe et al., 2016) |
| | 0.63 $^{(ii)}$ | | (Little et al., 1999) |
| | $0.2 - 0.4$ $^{(iii)}$ | | (Zeng et al., 2010) |
| $a$ | $2.7 \times 10^{-8}$ | $h^{-1} \,.\, cell^{-1}$ | (De Paepe & Taddei, 2006) |
| | $5 \times 10^{-8} - 3 \times 10^{-7}$ | | (De Paepe et al., 2016) |
| | $\sim 10^{-9} - 10^{-8}$ $^{(iv)}$ | | (Lindberg et al., 2014) |
| | $\sim 10^{-8} - 10^{-7}$ | | (Shao & Wang, 2008) |
| $B$ | 115 | $virus \,.\, cell^{-1}$ | (De Paepe & Taddei, 2006) |
| | 12.1 | | (De Paepe et al., 2016) |
| | $37 - 590$ $^{(iv)}$ | | (Lindberg et al., 2014) |
| | 56 | | (Little et al., 1999) |
| | $9.7 - 255$ | | (Wang, 2006) |
| $1/\tau$ | 0.7 | $h$ | (De Paepe & Taddei, 2006) |
| | $> 0.67$ | | (De Paepe et al., 2016) |
| | $0.78 - 1.67$ $^{(iv)}$ | | (Lindberg et al., 2014) |
| | 0.49-1.13 | | (Shao & Wang, 2008) |
| | 0.47-1.05 | | (Wang, 2006) |

$^{(i)}$ Virulent mutant $\lambda$cI*; $^{(ii)}$ MOI=6-8; $^{(iii)}$ MOI=1; $^{(iv)}$ Phages of *P. aeruginosa* from environmental water source.
