## Supplementary material for "Evolution of Virulence in Emerging Epidemics: From Theory to Experimental Evolution and Back": SI Appendix

July 26, 2024

**S1 A model coupling epidemiology and evolution** **2**

#### S1 A model coupling epidemiology and evolution

In the main text, we model in continuous cultures of *E. coli* the competition between two strains of phage $\lambda$: the wildtype strain – hereafter denoted $w$ – vs. the mutant strain (or variant) $\lambda$cI857 – hereafter denoted $m$. The variant $m$ is known to be more virulent than the wildtype strain $w$ due to a point mutation in the transcriptional repressor protein cI (St-Pierre & Endy, 2008; Sussman & Jacob, 1962).

##### S1.1 Epidemiology

We recall here the epidemiological model (system of ODEs) we use in the main text (see **Table 1** for notations):

$$\begin{cases} \dot{S}(t) &= rS(t)\left(1-\frac{N(t)}{K}\right)-\Big(abV(t)+\delta\Big)S(t) \\ \dot{L}(t) &= rL(t)\left(1-\frac{N(t)}{K}\right)+\overline{\phi}(t)abV(t)S(t)-\Big(\overline{\alpha}(t)+\delta\Big)L(t) \\ \dot{Y}(t) &= \big(1-\overline{\phi}(t)\big)abV(t)S(t)+\overline{\alpha}(t)L(t)-\Big(\tau+\delta\Big)Y(t) \\ \dot{V}(t) &= \tau Y(t)B-\Big(aN(t)+\delta\Big)V(t) \end{cases} \tag{S1}$$

where the overlines refer to mean values of the life-history traits after averaging over the distribution of strain frequencies:

$$\begin{cases} \overline{\alpha}(t) &= p(t)\alpha_m+\big(1-p(t)\big)\alpha_w \\ \overline{\phi}(t) &= q(t)\phi_m+\big(1-q(t)\big)\phi_w \end{cases} \tag{S2}$$

Furthermore, at each time point $t$, the prevalence is given by:

$$P(t) = \frac{Y(t)+L(t)}{N(t)} = 1-\frac{S(t)}{N(t)}, \tag{S3}$$

and its differentiation with respect to time yields:

$$\dot{P}(t)=\big(1-P(t)\big)\left[abV(t)-\left(r\left(1-\frac{N(t)}{K}\right)+\tau\right)\frac{Y(t)}{N(t)}\right]. \tag{S4}$$

##### S1.2 Evolution

We refer in the main text to the different frequencies associated with the mutant strain as follows:

- $p(t) = L_m(t)/L(t)$, the frequency of $L$ cells infected by the mutant strain;
- $q(t) = V_m(t)/V(t)$, the frequency of the mutant strain in the free virus stage ($V$);
- $f(t) = Y_m(t)/Y(t)$, the frequency of $Y$ cells infected by the mutant strain;
- $g(t) = \dfrac{Y_m(t) + L_m(t)}{Y(t) + L(t)}$, the frequency of cells infected (either $Y$ or $L$) by the mutant strain.

Note that we also have:

$$g(t) = f(t)\left(\frac{Y(t)}{Y(t)+L(t)}\right) + p(t)\left(\frac{L(t)}{Y(t)+L(t)}\right). \tag{S5}$$

Using our model (S1), we can then easily calculate the temporal dynamics of each of these frequencies, which yields:

$$\begin{cases} \dot{p}(t) = & \Big(\big(q(t)-p(t)\big)\phi_w + q(t)\big(1-p(t)\big)\Delta\phi\Big)\dfrac{abV(t)S(t)}{L(t)} - p(t)\big(1-p(t)\big)\Delta\alpha \\ \dot{q}(t) = & \Big(f(t)-q(t)\Big)\dfrac{Y(t)}{V(t)}\tau B \\ \dot{f}(t) = & \Big(\big(q(t)-f(t)\big)\big(1-\phi_w\big) - q(t)\big(1-f(t)\big)\Delta\phi\Big)\dfrac{abV(t)S(t)}{Y(t)} + \\ & \Big(\big(p(t)-f(t)\big)\alpha_w + p(t)\big(1-f(t)\big)\Delta\alpha\Big)\dfrac{L(t)}{Y(t)} \\ \dot{g}(t) = & \Big(p(t)-g(t)\Big)\left(r\left(1-\dfrac{N(t)}{K}\right)-\delta\right)\dfrac{L(t)}{Y(t)+L(t)} + \Big(q(t)-g(t)\Big)\dfrac{abV(t)S(t)}{Y(t)+L(t)} - \\ & \Big(f(t)-g(t)\Big)\big(\delta+\tau\big)\dfrac{Y(t)}{Y(t)+L(t)} \end{cases} \tag{S6}$$

with $\Delta\alpha = \alpha_m - \alpha_w$ and $\Delta\phi = \phi_m - \phi_w$.

Taken together, equations (S1)-(S6) yield the coupled evolutionary-epidemiological dynamics of this phage-bacteria system. Focusing instead on logit-frequencies, that is the log odds ln(frequency of the variant / frequency of the wildtype):

$$
\begin{cases}
\dfrac{\mathrm{d}\,\mathrm{logit}(p(t))}{\mathrm{d}t} & = \left(\dfrac{q(t)-p(t)}{p(t)\big(1-p(t)\big)}\phi_w + \dfrac{q(t)}{p(t)}\Delta\phi\right)\dfrac{abV(t)S(t)}{L(t)} - \Delta\alpha \\
\dfrac{\mathrm{d}\,\mathrm{logit}(q(t))}{\mathrm{d}t} & = \dfrac{f(t)-q(t)}{q(t)\big(1-q(t)\big)}\dfrac{Y(t)}{V(t)}\tau B \\
\dfrac{\mathrm{d}\,\mathrm{logit}(f(t))}{\mathrm{d}t} & = \left(\dfrac{q(t)-f(t)}{f(t)\big(1-f(t)\big)}(1-\phi_w) - \dfrac{q(t)}{f(t)}\Delta\phi\right)\dfrac{abV(t)S(t)}{Y(t)} + \\
& \quad \left(\dfrac{p(t)-f(t)}{f(t)\big(1-f(t)\big)}\alpha_w + \dfrac{p(t)}{f(t)}\Delta\alpha\right)\dfrac{L(t)}{Y(t)} \\
\dfrac{\mathrm{d}\,\mathrm{logit}\,(g(t))}{\mathrm{d}t} & = \dfrac{p(t)-g(t)}{g(t)\big(1-g(t)\big)}\left(r\left(1-\dfrac{N(t)}{K}\right)-\delta\right)\dfrac{L(t)}{Y(t)+L(t)} + \dfrac{q(t)-g(t)}{g(t)\big(1-g(t)\big)}\dfrac{abV(t)S(t)}{Y(t)+L(t)} - \\
& \quad \dfrac{f(t)-g(t)}{g(t)\big(1-g(t)\big)}\big(\delta+\tau\big)\dfrac{Y(t)}{Y(t)+L(t)}
\end{cases}
\tag{S7}
$$

### S2 Theoretical analyses

#### S2.1 Equilibrium states & basic reproduction number

In the absence of the virus, (S1) converges trivially to the following equilibrium state: $\big(S(\infty),\ L(\infty),\ Y(\infty),\ V(\infty)\big) = \big(K(1-\delta/r),\ 0,\ 0,\ 0\big)$ if $r > \delta$, $\big(0,\ 0,\ 0,\ 0\big)$ otherwise. When a single strain of the virus (with phenotypes $\alpha$ and $\phi$) is introduced in the bacterial population (fully susceptible, with density $S_0$), the fate of this phage-bacteria system depends on the basic reproduction number $\mathscr{R}_0$ of the pathogen – i.e., the expected number of secondary infections caused by one primary infected bacteria in an otherwise fully susceptible population. Using the next-generation-matrix method (Diekmann et al., 2010), we decompose the life cycle of phage $\lambda$ into transmission (matrix $\mathbf{T}$) and transition (matrix $\mathbf{\Sigma}$) components:

$$
\mathbf{T} = \begin{pmatrix} r\left(1-\frac{S_0}{K}\right) & 0 & \phi abS_0 \\ 0 & 0 & (1-\phi)abS_0 \\ 0 & 0 & 0 \end{pmatrix} \quad \text{and} \quad \mathbf{\Sigma} = \begin{pmatrix} -(\alpha+\delta) & 0 & 0 \\ \alpha & -(\tau+\delta) & 0 \\ 0 & \tau B & -(aS_0+\delta) \end{pmatrix},
$$

such that :

$$
\begin{pmatrix} \dot{L}(t) & \dot{Y}(t) & \dot{V}(t) \end{pmatrix}^\top = (\mathbf{T}+\mathbf{\Sigma})\begin{pmatrix} L(t) & Y(t) & V(t) \end{pmatrix}^\top
$$

The basic reproduction number $\mathscr{R}_0$ is given by the spectral radius of the matrix $-\mathbf{T}\mathbf{\Sigma}^{-1}$, that is:

$$\mathscr{R}_0 = \frac{A + \sqrt{A^2 - 4r\left(1 - \frac{S_0}{K}\right)(1-\phi)abS_0\tau B(aS_0+\delta)(\alpha+\delta)(\tau+\delta)}}{2(aS_0+\delta)(\alpha+\delta)(\tau+\delta)}, \tag{S8}$$

with $A = r\left(1 - \frac{S_0}{K}\right)(aS_0+\delta)(\tau+\delta) + abS_0\tau B(\alpha + (1-\phi)\delta)$. Note that, if $S_0 = K$, equation (S8) reduces to:

$$\mathscr{R}_0 = \frac{abK\tau B(\alpha + (1-\phi)\delta)}{(aK+\delta)(\alpha+\delta)(\tau+\delta)},$$

When $\mathscr{R}_0 < 1$, phages goes extinct and the bacterial population converges to the virus-free equilibrium above. Alternatively, when $\mathscr{R}_0 > 1$, an epidemic breaks out and eventually stabilises to the following endemic equilibrium:

$$\begin{cases} S(\infty) &= 0 \\ L(\infty) &= \dfrac{K\Big(r - (\delta+\alpha)\Big)}{\delta+\alpha+\tau}\ \dfrac{\delta+\tau}{r} \\ Y(\infty) &= \dfrac{K\Big(r - (\delta+\alpha)\Big)}{\delta+\alpha+\tau}\ \dfrac{\alpha}{r} \\ V(\infty) &= \dfrac{K\Big(r - (\delta+\alpha)\Big)}{\delta+\alpha+\tau}\ \dfrac{B\ \alpha\ \tau}{aK\Big(r-(\delta+\alpha)\Big) + \delta\ r} \end{cases} \tag{S9}$$

#### S2.2 Viral dynamics in an emerging epidemic

In the epidemic case, susceptible hosts are initially highly abundant. To simplify, we investigate the case where the density of susceptible hosts remains constant over time, i.e., $\forall t,\ S(t) = S_0$. Starting with a bacterial population at carrying capacity $K$, we also assume that the population size remains constant over time, i.e., $N(t) = K$, leading to the following simplified epidemiological system:

$$\begin{cases} \dot{S}(t) &= 0 \\ \dot{L}(t) &= \overline{\phi}(t)abV(t)S_0 - \Big(\overline{\alpha}(t) + \delta\Big)L(t) \\ \dot{Y}(t) &= \big(1 - \overline{\phi}(t)\big)abV(t)S_0 + \overline{\alpha}(t)L(t) - \Big(\tau+\delta\Big)Y(t) \\ \dot{V}(t) &= \tau Y(t)B - \Big(aK+\delta\Big)V(t) \end{cases} \tag{S10}$$

To study the selection gradient $\mathscr{S}$ of the virulent phage – i.e., the rate at which it grows or declines in frequency on the logit scale –, we focus on compartment $Y$. As shown in (S7), the temporal dynamics of $\text{logit}\big(f(t)\big)$ depends on the ratio $V(t)/Y(t)$, whose differentiation with respect to time is given by the quadratic polynomial:

$$\frac{\mathrm{d}}{\mathrm{d}t}\left(\frac{V(t)}{Y(t)}\right) = -\big(1-\overline{\phi}(t)\big)abS_0\left(\frac{V(t)}{Y(t)}\right)^2 - \left(\overline{\alpha}(t)\frac{L(t)}{Y(t)} + aK - \tau\right)\frac{V(t)}{Y(t)} + \tau B. \tag{S11}$$

We now use an argument of separation of time scale (Rinaldi & Scheffer, 2000; Verhulst, 2007) under the assumption of weak selection: epidemiological dynamics, such as $L(t)$, $Y(t)$ and $V(t)$, are treated as fast variables while evolutionary dynamics – i.e., strain frequencies – are treated as slow variables because phenotypic differences between the wildtype and the virulent strain are assumed to be small. Setting the right-hand side of (S11) to 0, we obtain the positive solution:

$$\frac{V(t)}{Y(t)} \approx \frac{\tau - \overline{\alpha}(t)\frac{L(t)}{Y(t)} - aK + \sqrt{\left(\tau - \overline{\alpha}(t)\frac{L(t)}{Y(t)} - aK\right)^2 + 4\big(1-\overline{\phi}(t)\big)abS_0\tau B}}{2\big(1-\overline{\phi}(t)\big)abS_0}.$$

In the early state of the epidemic, the system is mainly governed by the lytic pathway, so we assume that $L(t)/Y(t) \approx 0$ (**Fig. S3-B**). Furthermore, under the assumption of weak selection, we substitute $\overline{\phi}(t)$ by $\phi_w$. We denote $Z$ this approximation of the ratio $V(t)/Y(t)$:

$$Z = \frac{\tau - aK + \sqrt{(\tau - aK)^2 + 4(1-\phi_w)abS_0\tau B}}{2(1-\phi_w)abS_0}. \tag{S12}$$

The temporal dynamics of $\mathrm{logit}\big(f(t)\big)$ becomes:

$$\frac{\mathrm{d}\,\mathrm{logit}(f(t))}{\mathrm{d}t} \approx \underbrace{\frac{\tau - aK + \sqrt{(\tau - aK)^2 + 4(1-\phi_w)abS_0\tau B}}{2(1-\phi_w)}}_{abS_0 Z}\left(\frac{q(t)-f(t)}{f(t)\big(1-f(t)\big)}\big(1-\phi_w\big) - \frac{q(t)}{f(t)}\Delta\phi\right). \tag{S13}$$

We now look at the term in brackets. Posing $D(t) = \frac{q(t)-f(t)}{f(t)\big(1-f(t)\big)}$, we obtain $\frac{q(t)}{f(t)} = 1 + D(t)\big(1 - f(t)\big)$ and:

$$\frac{\mathrm{d}D(t)}{\mathrm{d}t} = \frac{\mathrm{d}\,\mathrm{logit}(q(t))}{\mathrm{d}t}\frac{q(t)\big(1-q(t)\big)}{f(t)\big(1-f(t)\big)} - \frac{\mathrm{d}\,\mathrm{logit}(f(t))}{\mathrm{d}t}\Big(1 + D(t)\big(1-2f(t)\big)\Big) \tag{S14}$$

We set the right-hand side of (S14) to 0 (quasi-equilibrium) and only keep the solution for which the term $\mathcal{O}(0)$ is equal to 0. Again, under the assumption of weak selection, phenotypic differences are small and $\mathcal{O}(\varepsilon)$. A Taylor expansion for the selected solution about the neutral case - i.e., when both strains share the same phenotype – to first order then yields:

$$D(t) \approx \frac{abS_0\left(\frac{V(t)}{Y(t)}\right)^2 \Delta\phi}{\tau B + abS_0\left(\frac{V(t)}{Y(t)}\right)^2 \big(1-\phi_w\big)} + \mathcal{O}(\varepsilon^2). \tag{S15}$$

Plugging approximations (S12) and (S15) into (S13), we obtain after some rearrangements:

$$\frac{\mathrm{d}\,\mathrm{logit}(f(t))}{\mathrm{d}t} \approx -\Delta\phi\ abS_0 Z \left(\frac{\tau B + abS_0 Z^2 \big(1-f(t)\big)\Delta\phi}{\tau B + abS_0 Z^2\big(1-\phi_w\big)}\right). \tag{S16}$$

We use (S16) as an approximation of the selection gradient of the virulent phage $\mathscr{S}$. As $f(t)$ is here always between 0.5 and 1 at the beginning of the epidemic:

$$-\Delta\phi\ abS_0 Z\left(\frac{\tau B + abS_0 Z^2 \Delta\phi/2}{\tau B + abS_0 Z^2\big(1-\phi_w\big)}\right) \leq \underbrace{\frac{\mathrm{d}\,\mathrm{logit}(f(t))}{\mathrm{d}t}}_{\mathscr{S}} \leq -\Delta\phi\ \frac{abS_0 Z\tau B}{\tau B + abS_0 Z^2\big(1-\phi_w\big)}, \tag{S17}$$

which may provide good approximations to predict the trajectory of the logit-frequency of the virulent phage in both compartment $Y$ and $V$, while $S(t)$ does not vary over time (**Fig. S3-C**).

Or, using once again an assumption of weak selection, a Taylor expansion of (S16) to first order in $\Delta\phi$ leads to:

$$\underbrace{\frac{\mathrm{d}\,\mathrm{logit}(f(t))}{\mathrm{d}t}}_{\mathscr{S}} \approx -\Delta\phi\ \frac{abS_0 Z\tau B}{\tau B + abS_0 Z^2\big(1-\phi_w\big)} + \mathcal{O}(\varepsilon^2), \tag{S18}$$

which is the approximation we use in the main text when we propose that, at the beginning of the epidemic:

$$\mathscr{S} \propto -\Delta\phi. \tag{S19}$$

Therefore, the virulent phage ($\Delta\phi < 0$) is selected for at the beginning of the epidemic.

##### S2.3 Viral dynamics at the end of the epidemic

At the end of the epidemic, the prevalence is high and the pool of susceptible host is depleted, so that $P(t) \approx 1$ and $S(t) \approx 0$; the system (S1) reduces then to:

$$
\begin{cases}
\dot{S}(t) &= 0 \\
\dot{L}(t) &= rL(t)\left(1-\dfrac{N(t)}{K}\right) - \left(\overline{\alpha}(t)+\delta\right)L(t) \\
\dot{Y}(t) &= \overline{\alpha}(t)L(t) - \left(\tau+\delta\right)Y(t) \\
\dot{V}(t) &= \tau Y(t)B - \left(aN(t)+\delta\right)V(t)
\end{cases}
\tag{S20}
$$

This phage-bacteria system is now driven by the lysogenic pathway so that the density of $Y$ cells becomes negligible compared to the density of $L$ cells. Indeed, using (S9):

$$
\lim_{S(t)\,\to\,0} \frac{Y(t)}{L(t)} = \frac{\alpha_w}{\delta+\tau} \approx 0,
$$

as $\alpha_w \ll \tau$ (time elapsed between phage integration and reactivation being much longer than lysis). According to (S5), this also means that the frequency of the virulent strain among infected hosts is now almost entirely driven by its frequency in lysogenic cells solely, that is $g(t) \approx p(t)$.

Rewriting the system (S20) in matrix form for the strain $k$ ($k \in \{w,\ m\}$):

$$
\begin{pmatrix} \dot{L}_k(t) \\ \dot{Y}_k(t) \\ \dot{V}_k(t) \end{pmatrix} = \underbrace{\begin{pmatrix} r\left(1-\frac{N(t)}{K}\right) - (\delta+\alpha_k) & 0 & 0 \\ \alpha_k & -(\tau+\delta) & 0 \\ 0 & \tau B & -(aN(t)+\delta) \end{pmatrix}}_{\mathbf{R}_k} \begin{pmatrix} L_k(t) \\ Y_k(t) \\ V_k(t) \end{pmatrix}, \tag{S21}
$$

the dominant eigenvalue of the Jacobian $\mathbf{R}_k$ is equals to $r\left(1-\frac{N(t)}{K}\right) - (\delta+\alpha_k)$. In these conditions, one would expect the selection gradient $\mathscr{S}$ of the mutant strain to be given in each compartment by the difference in eigenvalues which yields:

$$
\begin{aligned}
\mathscr{S} &= \left(r\left(1-\frac{N(t)}{K}\right) - (\delta+\alpha_m)\right) - \left(r\left(1-\frac{N(t)}{K}\right) - (\delta+\alpha_w)\right) \\
&= -\alpha_m + \alpha_w \;=\; -\Delta\alpha.
\end{aligned}
\tag{S22}
$$

As a result, the virulent phage is counter-selected in the long-term ($\Delta\alpha > 0 \;\Leftrightarrow\; \mathscr{S} < 0$) and, in each compartment, decreases in frequency at a rate of $\mathscr{S}$ on the logit scale (**Fig. S1-B**). We can once again predict the future trajectory of the logit-frequency of the virulent that linearly declines with negative slope $\mathscr{S} = -\Delta\alpha$. Note that recovering this result for compartment $L$ is straightforward as one just needs to set $S(t)$ to 0 in $\mathrm{d}\,\mathrm{logit}(p(t))/\mathrm{d}t$ in (S7).

##### S2.4 Differentiation across compartments

We define the differentiation of the virulent strain between free phages ($V$) and prophages ($L$) as:

$$\mathcal{Q}^{VL}(t) = \frac{q(t)}{1-q(t)}\frac{1-p(t)}{p(t)}, \tag{S23}$$

such that: $\ln\big(\mathcal{Q}^{VL}(t)\big) = \text{logit}(q(t)) - \text{logit}(p(t))$. Note that we also have $\mathcal{Q}^{VL}(t) = \frac{V_m(t)}{V_w(t)}\frac{L_w(t)}{L_m(t)}$ and that 1 corresponds to no differentiation. Using the eigenvectors associated with the dominant eigenvalues of the Jacobian matrices $\mathbf{R}_w$ and $\mathbf{R}_m$ (cf. equation (S21)), the differentiation $\mathcal{Q}^{YL}(t)$ converges towards:

$$\mathcal{Q}^{VL} = \frac{\alpha_m\big(r(1-N(t)/K) - \alpha_w + \tau\big)}{\alpha_w\big(r(1-N(t)/K) - \alpha_m + \tau\big)} \approx \frac{\alpha_m}{\alpha_w} = 1 + \frac{\Delta\alpha}{\alpha_w}, \tag{S24}$$

as $\alpha_m$ and $\alpha_w$ are very small values.

Likewise, the differentiation between $Y$ and $L$ cells:

$$\mathcal{Q}^{YL}(t) = \frac{f(t)}{1-f(t)}\frac{1-p(t)}{p(t)}, \tag{S25}$$

converges towards:

$$\mathcal{Q}^{YL} = \frac{\alpha_m\big(r(1-N(t)/K) - \alpha_w + aN(t)\big)\big(r(1-N(t)/K) - \alpha_w + \tau\big)}{\alpha_w\big(r(1-N(t)/K) - \alpha_m + aN(t)\big)\big(r(1-N(t)/K) - \alpha_m + \tau\big)} \approx \frac{\alpha_m}{\alpha_w} = 1 + \frac{\Delta\alpha}{\alpha_w} \tag{S26}$$

and the differentiation between free phages ($V$) and $Y$ cells:

$$\mathcal{Q}^{VY}(t) = \frac{q(t)}{1-q(t)}\frac{1-f(t)}{f(t)} \tag{S27}$$

converges towards:

$$\mathcal{Q}^{VY} = \frac{r(1-N(t)/K) + aN(t) - \alpha_w}{r(1-N(t)/K) + aN(t) - \alpha_m} \approx 1. \tag{S28}$$

When the system stabilizes (endemic or late stage of the epidemic case), the virulent strain is therefore more frequent among free viruses and $Y$ cells than among $L$ cells (prophages) but we also expect almost no differentiation between free viruses and $Y$ cells (**Fig. S4**).

#### S3 Statistical inference of the rates of prophage reactivation

From the previous analysis of the model, we show that when the system reaches high prevalence the selection gradient $\mathscr{S}$ of the variant is simply given by: $\mathscr{S} = \alpha_w - \alpha_m = -\Delta\alpha$ (S22); and the differentiation $\mathcal{Q}^{VL}$ of the variant between free phages and prophages by: $\mathcal{Q}^{VL} = \alpha_m/\alpha_w = 1 + \Delta\alpha/\alpha_w$ (S24).

Three quantities are tracked over time in the experiment: (i) the prevalence $P(t)$, (ii) the frequency of hosts infected by the virulent phage $g(t)$ and (iii) the frequency of the virulent phage in the free virus stage $q(t)$. We only keep data from the time point the system has reached high prevalence ($\geq 95\%$). We recall that, in these conditions, $g(t)$ is almost entirely driven by the frequency of $L$ cells infected by the virulent phage, that is $g(t) \approx p(t)$ (cf. **§S2.3**).

##### S3.1 Frequentist approach

We fit a linear mixed-effects model on the logit-frequency infected by the virulent phage $\operatorname{logit}(g(t))$ to estimate the slope $\mathscr{S}$. For treatment $i$ (epidemic vs. endemic), in chemostat $j$ and at time point $t$, we have ($t$, $i$ and $j$ are now noted as indexes for clarity):

$$\underbrace{\operatorname{logit}(g)_{i,j,t}}_{\text{Response variable}} = intercept + \mathscr{S} \times t + \beta_i + \chi_j + \varepsilon_{i,j,t},$$

with:

- $intercept$, the common fixed effect (reference);
- $\beta_i$, the fixed effect of treatment $i$ on the intercept of the model;
- $\chi_j \overset{i.i.d}{\sim} \mathcal{N}(0,\ \gamma^2)$, the random effect (with variance $\gamma^2$) of the $j^{\text{th}}$ chemostat on the intercept of the model;
- $\varepsilon_{i,j,t} \overset{i.i.d}{\sim} \mathcal{N}(0,\ \sigma_g^2)$, the residual error (with variance $\sigma_g^2$).

We fit this mixed-effects model using the function '*lmer*' from the R package '*lme4*'. Alongside, we also propose a version where the rates of prophage reactivation $\alpha_w$ and $\alpha_m$ (and therefore the slope $\mathscr{S}$) are allowed to vary across chemostats; this slightly changes the previous linear model to:

$$\operatorname{logit}(g)_{i,j,t} = intercept + (\mathscr{S} + \kappa_j) \times t + \beta_i + \chi_j + \varepsilon_{i,j,t},$$

with $\kappa_j$ the fixed effect of the $j^{\text{th}}$ chemostat on the slope of the model.

In parallel, we estimate $\mathcal{Q}^{VL}$, starting from the computation of $\ln\left(\mathcal{Q}^{VL}(t)\right) = \text{logit}\left(q(t)\right) - \text{logit}\left(p(t)\right)$, in which we substitute $\text{logit}\left(p(t)\right)$ by $\text{logit}\left(g(t)\right)$. Using the same notations for the indexes we have:

$$\underbrace{\text{logit}\left(q\right)_{i,j,t} - \text{logit}\left(g\right)_{i,j,t}}_{\ln(\mathcal{Q}^{VL})_{i,j,t}\text{, response variable}} = \ln\left(\mathcal{Q}^{VL}\right) + \varepsilon_{i,j,t},$$

where $\varepsilon \overset{i.i.d}{\sim} \mathcal{N}\left(0,\ \sigma_Q^2\right)$ is the residual error (with variance $\sigma_Q^2$). We thus estimate $\mathcal{Q}^{VL}$ by computing, all chemostats combined, the exponential of the arithmetic mean of $\ln\left(\mathcal{Q}^{VL}(t)\right)$ – this is completely equivalent to computing the geometric mean of $\mathcal{Q}^{VL}(t)$.

Alternatively, if we want to allow $\alpha_w$ and $\alpha_m$ (and therefore $\mathcal{Q}^{VL}$) to vary across chemostats:

$$\text{logit}\left(q\right)_{i,j,t} - \text{logit}\left(g\right)_{i,j,t} = \ln\left(\mathcal{Q}^{VL}\right)_j + \varepsilon_{i,j,t},$$

and we just need to compute the mean for each chemostat.

Finally, we obtain point estimates of both rates of prophage reactivation $\alpha_w$ and $\alpha_m$ combining equations (S22) and (S24):

$$\begin{cases} \alpha_w &= \dfrac{\mathscr{S}}{1-\mathcal{Q}^{VL}} \\ \alpha_m &= \dfrac{\mathscr{S}\times\mathcal{Q}^{VL}}{1-\mathcal{Q}^{VL}} \end{cases} \tag{S29}$$

in which we use the estimated values of $\mathscr{S}$ and $\mathcal{Q}^{VL}$. From our experimental data, all chemostats combined, we get (expressed in $h^{-1}$): $\hat{\alpha}_w = 2.58\times10^{-3}$ and $\hat{\alpha}_m = 1.19\times10^{-2}$.

#### S3.2 Bayesian approach

Besides, we also use a Bayesian approach to estimate parameters $\alpha_w$ and $\alpha_m$ along with their 95% credible intervals. For treatment $i$ (epidemic vs. endemic), in chemostat $j$ and at time point $t$, we consider the following likelihoods for the response variables ($t$, $i$ and $j$ are now noted as indexes for clarity):

$$\begin{cases} \text{logit}\left(g\right)_{i,j,t} \sim \mathcal{N}\Big(\beta_i + \chi_j + \overbrace{(\alpha_w-\alpha_m)}^{\mathscr{S}}\times t,\ \sigma_g^2\Big) \\ \Big(\text{logit}\left(q\right)_{i,j,t} - \text{logit}\left(g\right)_{i,j,t}\Big) \sim \mathcal{N}\Big(\underbrace{\ln(\alpha_m)-\ln(\alpha_w)}_{\ln(\mathcal{Q}^{VL})},\ \sigma_Q^2\Big) \end{cases}$$

with, as before:

- $\beta_i$, the fixed effect of treatment $i$ on the intercept of the model;

- $\chi_j \sim \mathcal{N}\left(0,\ \gamma^2\right)$, the random effect (with variance $\gamma^2$) of the $j^{\text{th}}$ chemostat on the intercept of the model;

- $\sigma_g^2$ and $\sigma_Q^2$, the variance parameters of the response variables.

We choose the following prior distributions:

$$\begin{aligned}
\alpha_w &\sim \mathcal{U}\left([0,\ 0.1]\right)\\
\alpha_m &\sim \mathcal{U}\left([0,\ 0.1]\right)\\
\beta_{epidemic} &\sim \mathcal{U}\left([0.25,\ 3]\right)\\
\beta_{endemic} &\sim \mathcal{U}\left([-0.25,\ 0.25]\right)\\
\gamma &\sim \mathcal{U}\left([0,\ 3]\right)\\
\sigma_Q &\sim \mathcal{U}\left([0,\ 5]\right)\\
\sigma_g &\sim \mathcal{U}\left([0,\ 1]\right)
\end{aligned}$$

Again, if we allow parameters $\alpha_w$ and $\alpha_m$ to vary across chemostats:

$$\begin{cases}
\text{logit}\left(g\right)_{i,j,t} \sim \mathcal{N}\Big(\beta_i + \chi_j + (\alpha_{w,j} - \alpha_{m,j}) \times t,\ \sigma_g^2\Big)\\
\Big(\text{logit}\left(q\right)_{i,j,t} - \text{logit}\left(g\right)_{i,j,t}\Big) \sim \mathcal{N}\Big(\ln(\alpha_{m,j}) - \ln(\alpha_{w,j}),\ \sigma_Q^2\Big)
\end{cases}$$

where the prior of each $\alpha_{w,j}$ and $\alpha_{m,j}$ is the same as for $\alpha_w$ and $\alpha_m$ above.

We independently run 4 Monte-Carlo Markov chains using JAGS (Plummer et al., 2003) version 4.3.0 and function '*jags*' from the R package '*R2jags*'. Each chain is 20,000 iterations long (length of the burn-in period 8,000) with thinning rate 50. As initial conditions: $\alpha_w$ and $\alpha_m$ are drawn uniformly between 0 and $3 \times 10^{-2}$ such that $\alpha_w < \alpha_m$; $\beta_{epidemic}$ is drawn uniformly between 0.25 and 3, and $\beta_{endemic}$ between -0.25 and 0.25; $\chi_j$ is drawn from a standard Normal distribution; and variance parameters $\sigma_Q$, $\sigma_g$ and $\gamma$ are drawn uniformly between 0 and 1. In the end, we assess convergence of posterior distributions, especially we check that Gelman-Rubin statistics are below 1.1 and that effective sample sizes are above 100.

From our experimental data, all chemostats combined, we get (expressed in $h^{-1}$): $\alpha_w = 2.65 \times 10^{-3}$ (mean, 95% credible interval $[2.03 \times 10^{-3},\ 3.32 \times 10^{-3}]$) and $\alpha_m = 1.21 \times 10^{-2}$ (mean, 95% credible interval $[9.48 \times 10^{-3},\ 1.47 \times 10^{-2}]$) (**Fig. S14-A**); see **Fig. S14-B** for posterior distributions by chemostat.
